# Leaflet shape modulates canopy light-use efficiency by partitioning light more evenly among soybean leaves

**DOI:** 10.64898/2026.11.20.768164

**Authors:** Bishal G. Tamang, Anthony Digrado, Carl J. Bernacchi, Elizabeth A. Ainsworth

**Affiliations:** Carl R. Woese Institute for Genomic Biology, University of Illinois Urbana-Champaign, Urbana, IL 61801, USA; Department of Plant Biology, University of Illinois Urbana-Champaign, Urbana, IL 61801, USA; Department of Crop Sciences, University of Illinois Urbana-Champaign, Urbana, IL 61801, USA

**Keywords:** canopy architecture, canopy photosynthesis, crop modeling, *Glycine max*, *GmJAG1*, leaf area index, three-dimensional ray tracing

## Abstract

- Modern soybean (*Glycine max*) canopies carry more leaves than they need, so yield gains depend on using light more efficiently. Lines carrying the narrow-leaflet allele of *GmJAG1* yield as much as broad-leaflet lines despite having less leaf area. We asked whether leaflet shape raises canopy light capture or light-use efficiency.
- To separate the two, we simulated 530 measured plots of broad- and narrow-leaflet near-isogenic lines at two sites in the Helios three-dimensional ray-tracing model. We compared these simulations with a one-dimensional model of identical leaf physiology and with field data, then mapped the traits by genome-wide association.
- In the simulations, both leaflet types absorbed similar fractions of light, but narrow-leaflet canopies had 4-7% higher light-use efficiency. The one-dimensional model recovered little to none of this gain, tying it to canopy structure, where narrow leaflets partitioned light more evenly among leaves at each depth. In the field, narrow lines matched broad-line yield with 12% less leaf area, and yield per unit leaf area (+13.7%) mapped to *GmJAG1*, as did leaflet shape and simulated traits.
- With less leaf area, narrow-leaflets canopies are thus predicted to absorb as much light as broad-leaflet canopies and use it more efficiently. As a heritable single-gene trait, leaflet shape could complement leaf-biochemical routes to improved photosynthesis.

## Introduction

The yield potential (Y_p_) of crops is the product of incident solar radiation and the efficiencies of light interception (ε_i_), the conversion of intercepted radiation into biomass (ε_c_), and the partitioning of biomass to harvested product (ε_p_) (Monteith, 1977; Zhu et al., 2010). Among these, breeding and agronomic management have brought light-interception efficiency and harvest index close to their theoretical maxima in major grain crops, leaving conversion efficiency (ε_c_, primarily determined by photosynthesis) as the remaining target for further gains in yield potential (Long et al., 2006; Zhu et al., 2010). Canopy-level ε_c_ depends on both the photosynthetic capacity of individual leaves and the spatial distribution of light within the canopy, with the latter shaped by canopy architecture (Yin & Struik, 2015; Slattery & Ort, 2021). Yet, both theoretical and empirical analyses have consistently shown that crop canopies overinvest in light interception, leading to poor light utilization (Zhu et al., 2010; Srinivasan et al., 2017; Slattery & Ort, 2021). Canopy architecture is therefore a target for raising ε_c,_ alongside leaf biochemistry.

Canopy architecture acts on photosynthesis through the light each leaf receives. Leaf size, leaf shape, leaf angle, leaf arrangement and internode length together set the within-canopy light environment. This environment in turn determines whether leaves at any given canopy depth operate under light-saturation where absorbed light exceeds photosynthetic capacity, in the efficient light-limited (near-linear) region, or below the light-compensation point (Niinemets, 2010; Niinemets et al., 2015; Slattery & Ort, 2021). Consistent with this, functional-structural plant models have demonstrated that small changes in architectural traits can produce measurable canopy photosynthesis effects. For instance, in a simulation of tomato plants, increasing the leaf length-to-width ratio by 1.5-fold increased canopy light absorption by 8-10% and photosynthesis by ∼6% (Sarlikioti et al., 2011). At a larger scale, a 3D ray-tracing simulation of soybean canopies attributed ∼17% of the elevated-CO_2_ gross primary productivity gain to canopy structural changes (primarily increased leaf area) (Song et al., 2020). At a global scale, satellite-derived canopy clumping architecture explained substantial across-region variation in crop yield over the 2001-2016 record across the four major staple crops, with more clumped canopy architectures (up to an intermediate optimum clumping index of 0.55-0.58) consistently associated with higher yields and lower nitrous oxide emissions (Yan et al., 2026). These results indicate that architectural traits can be engineered to increase productivity, but such predictions need testing in field canopies that differ in a single, defined architectural trait.

Soybean leaflet shape offers such a test. Leaflet shape is under simple genetic control by the *Ln* locus (*GmJAG1*) on chromosome 20, which encodes a transcription factor and distinguishes broad-leaflet from narrow-leaflet phenotypes through a single amino acid substitution in the EAR repression motif (Jeong et al., 2012; Fang et al., 2013). The narrow-leaflet allele also pleiotropically increases the number of seeds per pod (Jeong et al., 2012; Sayama et al., 2017). In addition, lanceolate leaves are anatomically thicker than ovate leaves, with approximately 30 μm of additional thickness per unit increase in leaf length-to-width ratio, a higher proportion of spongy mesophyll, and higher area-based photosynthesis and stomatal conductance (Tamang et al., 2023). Despite these leaf-level differences and contrasting canopy structure, comparative agronomic studies of narrow-and broad-leaflet lines have generally reported similar yield for the two leaflet types (Mandl & Buss, 1981; Wells et al., 1993; Dinkins et al., 2002). By contrast, CRISPR knockout of *GmJAG1* and its homeolog *GmJAG2* in a low-latitude soybean cultivar produced an ∼8.7% yield increase across two field trials (Cai et al., 2021), indicating that benefits of the narrow-leaflet trait can be context-dependent.

Recent work has begun to address the canopy-scale mechanism underlying the narrow-leaflet trait. Bianchi et al. (2020) compared two pairs of lanceolate and ovate near-isogenic lines and reported higher intercepted photosynthetically active radiation (PAR) per unit leaf area, higher net assimilation rate, and elevated red-to-far-red ratios at canopy mid-strata in lanceolate canopies. They proposed photomorphogenic signaling as the canopy-level mechanism linking leaflet morphology to reproductive output. Zhao et al. (2023) then measured canopy apparent photosynthesis directly in lanceolate and ovate cultivars under contrasting density and nitrogen treatments. They reported higher canopy light-and nitrogen-use-efficiency in lanceolate canopies, with the lanceolate advantage tracing predominantly to greater photosynthetic utilization of the direct light component. Most recently, Tamang et al. (2026) used near-isogenic lines across two locations and two row spacings to demonstrate that narrow-leaflet lines produced equivalent yield to their broad-leaflet counterparts (5,756 vs 5,801 kg ha^-1^, P = 0.43) despite 13% lower peak leaf area index and 3% lower digital biomass. That study, however, did not measure or model canopy carbon gain. These convergent findings indicate that narrow-leaflet canopies operate at higher canopy efficiency. Explaining how narrow lines yield as much from less leaf area requires separating two routes, light interception and light-use efficiency (LUE), at plot scale in a near-isogenic background.

Decomposing canopy net photosynthesis (canopy A) into absorbed PAR and light-use efficiency requires a model that resolves the spatial distribution of light within a structurally heterogeneous canopy. Big-leaf and two-leaf canopy models that integrate sun-shade fractions without resolving within-canopy heterogeneity capture some architectural effects, but they cannot distinguish leaflet-shape effects on the within-canopy light field from effects of total leaf area (de Pury & Farquhar, 1997; Gu et al., 2022). For example, a recent comparison in maize reported that crop yield predictions decreased by 12 to 48.5% with increasing dimensional resolution of canopy light simulation, and that 0D, 1D and 2D canopy models failed to capture inter-cultivar architectural differences resolved by a 3D representation (Gu et al., 2022). Resolving the leaflet-shape mechanism in soybean therefore requires a 3D representation of canopy structure with explicit ray-tracing of within-canopy light.

Here, we used the Helios 3D ray-tracing canopy biophysics framework (Bailey, 2019), parameterized with field-measured per-plot leaf area index, plant height, leaflet morphology, and leaf-level biochemistry from 200 BC_3_F_2:4_ near-isogenic lines (530 plots) at two sites and two row spacings (38 cm and 76 cm). Our objectives were (i) to test whether soybean leaflet shape modulates canopy carbon gain via light interception or via light-use efficiency, (ii) to identify where in the canopy the leaf-shape effect operates, (iii) to quantify the contribution of 3D canopy structure that lower-dimensional abstractions cannot capture, and (iv) to test whether per-line variation in the predicted canopy outputs traces to the *GmJAG1* leaflet-shape locus. Narrow and broad lines absorbed similar fractions of light, but narrow canopies were predicted to use it more efficiently by spreading it more evenly among leaves at each depth. A one-dimensional model with the same leaf physiology missed this gain, which therefore came from canopy structure and mapped, with leaflet shape and yield per leaf area, to *GmJAG1*. Leaflet shape is thus a single-gene canopy trait that raises predicted light-use efficiency and lets narrow lines yield as much as broad lines from less leaf area.

## Materials and Methods

### Plant material and experimental design

We studied 200 BC_3_F_2:4_ near-isogenic lines (NILs) of soybean differing at the *Ln* locus controlling leaflet shape. The recurrent parent was LD11-2170 (Maturity Group III), an elite broad-leaflet line developed at the University of Illinois and used as the reference for Group III in the uniform soybean yield tests in the Northern Region of the US (Cai & Brock, 2023). The narrow-leaflet donors were PI 612713A (MG I) and PI 547745 (MG II), both carrying the narrow-leaflet phenotype identified in a 2020 USDA germplasm survey (Tamang et al., 2023). The breeding history and yield characterization of this panel were reported by Tamang et al. (2026).

Field experiments were conducted during the 2024 growing season at two University of Illinois sites, SoyFACE (SF) and Energy Farm (EF), each at two row spacings (38 cm and 76 cm), giving four (location x row spacing) combinations (Fig 1A, B). Plots comprised four 2.4-m rows separated by 0.9-m alleys and were bordered on all sides by four rows of a commercial variety. Seeds were mechanically planted on 30 May 2024 (EF) and 4 June 2024 (SF) at 314,000 seeds ha^-^¹ (76 cm) and 447,000 seeds ha^-^¹ (38 cm). Plots followed a Type II Modified Augmented Design (MAD; Lin & Poushinsky, 1985), with 25 plots per block and LD11-2170 as the central primary check and the donor parents randomly distributed as secondary checks. SoyFACE comprised 9 blocks at 76 cm and 5 blocks at 38 cm while Energy Farm comprised 5 blocks at each spacing. In total, 606 plots were planted (542 NIL test plots and 64 parent check plots). Weeds were removed manually as needed, and irrigation was applied during early vegetative growth (once at SoyFACE, twice at Energy Farm).

**Figure 1.**
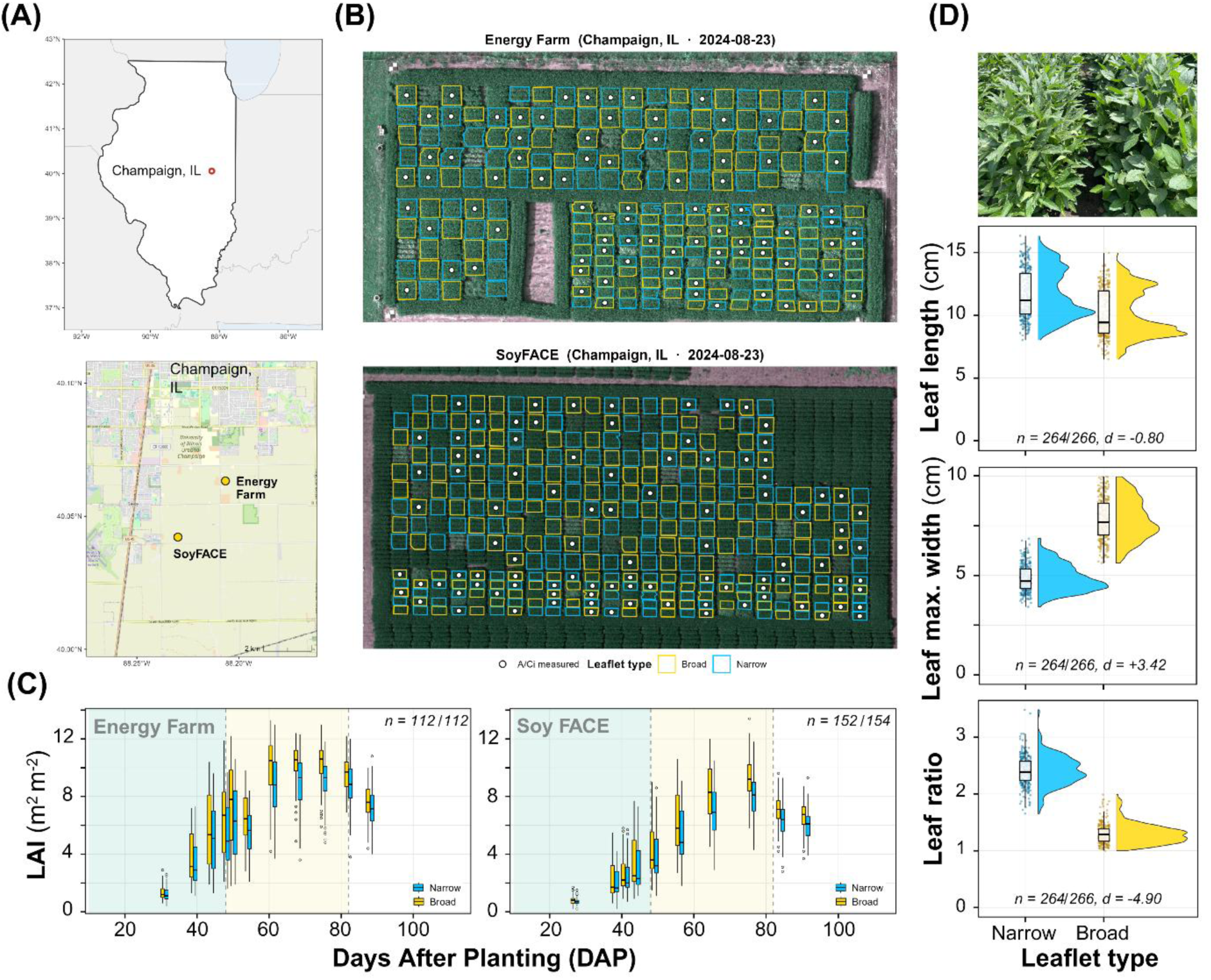
Field trials and ground-truth measurements of broad- and narrow-leaflet near-isogenic lines (NILs) of soybean at two Illinois sites in 2024 (n = 530 plots). **(A)** Locations of the SoyFACE and Energy Farm sites near Champaign, Illinois. **(B)** Aerial orthomosaics of both sites with plots outlined by leaflet type. Markers show plots where leaf gas-exchange (A-Ci) curves were measured. **(C)** Per-plot leaf area index (LAI) against days after planting (DAP) by leaflet type. Boxes show medians and interquartile ranges, and background shading marks the vegetative, reproductive and maturity stages. **(D)** Adjacent narrow- and broad-leaflet rows at canopy closure (top) and per-plot distributions of leaflet length, maximum width and length-to-maximum-width ratio (bottom). Points are plots, boxes show medians and interquartile ranges, and each panel gives n (narrow/broad plots) and Cohen’s d (broad minus narrow, pooled SD).

All analyses used the isogenic test plots, except the SoyFACE comparisons of light and leaf-area profiles between leaflet types, which also included six plots of the broad-leaflet recurrent parent LD11-2170 (pooled with the broad-leaflet NILs; Methods S1.8). Of the 542 NIL plots, 530 had complete leaf area index (LAI) time courses (SF n = 306; EF n = 224; plot list in Table S1), and 515 of these (SF n = 300; EF n = 215) had complete per-plot yield records. Leaf photosynthetic capacity was estimated from the response of net CO_2_ assimilation (*A*) to intercellular CO_2_ concentration (C_i_), measured by infrared gas exchange. The maximum carboxylation rate of Rubisco at 25 °C (Vcmax25) was estimated for 198 plots (SF n = 100; EF n = 98), and the maximum electron transport rate at 25 °C (Jmax25) for 164 of these (SF n = 86; EF n = 78). Where a per-plot biochemical or optical measurement was unavailable, the plot received the mean of its (location x row spacing x leaflet shape) combination. Per-trait sample sizes are reported in Fig S1.

### Field measurements

LAI was measured within each plot with a SunScan Canopy Analysis System (SS1; Delta-T Devices, Cambridge, UK) on 10 dates at SF (days after planting; DAP 27-91) and EF (DAP 31-88), recording one value per plot per date with the probe at the plant base, perpendicular to the rows. Plant height was measured manually at approximately the same schedule and interpolated to each LAI date (per-plot per-DAP height and LAI in Table S2). Leaflet length, average and maximum width and area were measured with an LI-3000C portable area meter (LI-COR BioSciences, Lincoln, NE, USA) on a fully expanded middle trifoliate between the 5th and 7th nodes at the V5-V7 stage (per-plot values in Table S3; per-plot distributions in Fig S2). Leaflet length-to-maximum-width ratio was calculated as length divided by maximum width.

Vcmax25 and Jmax25 were estimated from A-Ci response curves on excised, recently matured trifoliate leaves following established protocols (Long & Bernacchi, 2003; Busch et al., 2024; leaf sampling and handling in Methods S1). Gas exchange used LI-6800 systems (LI-COR Biosciences, Lincoln, NE, USA), each with a 6-cm² cuvette, and curves were fitted by the whole-curve method in PhotoGEA v1.1.0 (Lochocki et al., 2025), using the temperature responses of Bernacchi et al. (2001) (per-plot fitted parameters in Table S4). On the same leaf, chlorophyll index (mean of four SPAD-502 readings; Konica Minolta, Tokyo, Japan), leaf mass per area (LMA; gravimetric on a subset of plots), and reflectance spectra (350-2500 nm at 1-nm resolution, mean of four scans per leaf; ASD FieldSpec with contact-probe leaf clip, ASD Inc., Boulder, CO, USA) were recorded. For plots without these same-leaf measurements, values from one leaf per plot measured in a survey on 7-9 August 2024 were used (number of plots in Figs S1 and S3).

Weather data (air temperature, relative humidity, wind speed, barometric pressure, incoming shortwave irradiance, and PAR) were logged at each site by a weather station. Air temperature, relative humidity, wind speed and barometric pressure, averaged over 11:00-15:00 CST on each LAI measurement date, were supplied as per-DAP atmospheric inputs to the simulations (per-site per-DAP values in Table S5; daily aggregates in Fig S4; barometric pressure in Methods S1). Plots were harvested at maturity with an ALMACO R1 single-plot rotary combine (ALMACO, Nevada, IA, USA), which recorded plot weight and grain moisture (central two rows at 76 cm, all four rows at 38 cm; per-plot yield in Table S6).

For comparison with the simulated light profile, canopy PAR interception was measured at SoyFACE under 76-cm spacing on 20-22 August 2024 (DAP 77-79) with LI-190R quantum sensors (LI-COR) mounted at eight heights (5, 20, 40, 60, 80, 100, 120 and 140 cm) on a vertical pole, the Light Transmission Efficiency Scaffold (LITES), averaged over three scans per plot. The fraction of intercepted PAR (fIPAR) at the six within-canopy heights (5-100 cm) was computed relative to PAR at 140 cm (Methods S1.9). The value at the canopy floor (5 cm) gave whole-canopy interception. Because the sensors record only the PAR that reaches them, fIPAR counts PAR reflected back out of the canopy as well as PAR absorbed by leaves. Measured fIPAR was therefore expected to exceed the fraction of incident PAR absorbed (fAPAR) simulated by Helios. We report their difference, fIPAR – fAPAR (Methods S1; per-plot profiles in Table S7). During the same campaign, vertical canopy leaf-area profiles were obtained at these plots with the SS1 SunScan probe placed at the same six heights (5-100 cm) for two of the three pole positions, yielding cumulative LAI above each height (distinct from the routine plant-base LAI).

To map foliage distribution at the second site, canopy structure was measured from above at EF under 76-cm spacing with a 128-beam LiDAR sensor (Ouster OS0-128; Ouster Inc., San Francisco, CA, USA) mounted on a cable-suspended SpiderCam platform (Ross Video, Ottawa, Canada), using the same platform and point-cloud acquisition described in Tamang et al. (2026). For each plot, we derived 99th-percentile canopy height, the median relative height of foliage returns (“median foliage height”), and the share of foliage returns above 70% of canopy height (processing in Methods S1.11). Analysis used the 112 NIL plots at this site and spacing (56 Broad and 56 Narrow), the same plots simulated in Helios (plot metrics in Tables S8-S9).

### Per-plot leaf optical properties

Per-plot PAR (400-700 nm) and NIR (700-2500 nm) reflectance and transmittance were retrieved with PROSPECT-D (Féret et al., 2017) following the two-tier strategy of Wang et al. (2026). The two most variable biochemical parameters, leaf chlorophyll content (from SPAD via LCC = 1.26 x SPAD + 6.63) and LMA, were anchored per plot. The remaining structural and pigment parameters, which are harder to estimate per plot, were fitted per location x row spacing combination from that combination’s mean reflectance spectrum. Details are in Methods S1.3, with per-plot bands in Table S10, inversion parameters in Table S11 and leaf optical inputs in Fig S3.

### Three-dimensional canopy simulations

Canopy net photosynthesis at each plot and date was simulated in Helios, an open-source 3D ray-tracing canopy biophysics framework (Bailey, 2019; release v1.3.71). Parameterization followed Wang et al. (2026) for soybean in the same season and climate region except where noted. Each plot was simulated as a 3 x 3 m canopy with periodic horizontal boundaries and a planophile leaf-angle distribution. Canopy structure came from each plot’s field data on each date: leaflet length and average width, LAI and plant height. Leaves were placed at random within a horizontally homogeneous canopy, so row spacing entered the simulations only through each plot’s measured LAI and height. Each leaflet was divided into nine flat sub-leaf elements, the surfaces on which absorbed light and photosynthesis were computed. To test sensitivity to this choice, the peak LAI simulation for each location x row spacing x leaflet shape combination was repeated with 1, 4, 9, 25, 64, 144 and 225 elements per leaflet (Table S12). Leaf PAR/NIR optics were the per-plot PROSPECT-D values and longwave emissivity was 0.95. Solar position was computed at each site at 12:00 CDT from site coordinates (SF 40.033 °N, 88.233 °W; EF 40.063 °N, 88.207 °W), and direct and diffuse sunlight for a clear sky were computed with the REST2 model (Gueymard, 2008). Soil reflectivity, the number of light-scattering iterations, aerosol turbidity and ambient CO_2_ (415 ppm) are listed in Table S13.

Within-canopy Vcmax25 followed the exponential profile used by Wang et al. (2026), Vcmax25(l) = Vcmax25(top) x exp (0.64 l), where l is relative canopy height (−1 at base, 0 at top). Vcmax25(top) was set per plot from the field-measured leaf value. Jmax25, by contrast, was held uniform at the value measured at the top of the canopy. Dark respiration (Rd25), from the same PhotoGEA fits, was averaged within each (location x row spacing x leaflet shape) combination, giving eight combination-mean values (Table S14) applied uniformly within each combination. Temperature dependencies of the Farquhar-von Caemmerer-Berry (FvCB) photosynthesis model (Farquhar et al., 1980) followed Bernacchi et al. (2001), with Jmax activation energy corrected per Sharkey et al. (2007). The full constant set and the coupled stomatal-energy-balance solution are detailed in Methods S1. Absorbed PAR was computed by reverse ray-tracing (Bailey, 2018).

Helios was run for all 530 plots on each measurement date (5,299 plot-date combinations) on NVIDIA A40 graphics processors. Each plot used its own Vcmax25, Jmax25 and triose phosphate utilization (Tp25) where these were measured and the combination mean otherwise. For each plot and date, the simulation returned canopy A, fAPAR and canopy LUE (canopy A divided by absorbed PAR; Table S15). Absorbed PAR of each sub-leaf element was also used to describe how absorbed PAR was distributed with depth and among elements at the same depth (Methods S1.10).

### One-dimensional sun-shade model comparison

To isolate the contribution of 3D canopy structure, Helios outputs were compared against a 1D sun-shade two-leaf model following de Pury & Farquhar (1997). The model used the same leaf biochemistry, within-canopy profiles, temperature responses, leaf energy-balance and stomatal models, and NIR optics as Helios. The 1D model differed mainly in radiation transport, using a ten-layer two-stream scheme with the direct/diffuse partitioning of Spitters et al. (1986) instead of 3D ray tracing (Methods S1.6). The architectural residual was the narrow-minus-broad difference in canopy A predicted by Helios minus that predicted by the 1D model, computed within each location x row spacing combination (per-plot per-DAP 1D canopy A in Table S16). To check that filling unmeasured plots with combination means did not drive the residual, we repeated the comparison using only the 164 plots with both Vcmax25 and Jmax25 measured (Methods S1).

### Statistical analysis

Broad-versus-narrow differences in Helios-predicted canopy net photosynthesis, fAPAR, and canopy light-use efficiency were tested within each location x row spacing combination by two-sided Wilcoxon rank-sum tests on per-plot seasonal peak values, with Cohen’s *d* effect sizes (pooled SD; Cohen, 1988; positive = Broad > Narrow). Equivalence of post-closure fAPAR between leaflet types was tested with two one-sided Welch t-tests (TOST) against a ±0.02 margin. Per-plot yield, peak LAI and yield per unit peak LAI were compared by two-sided Welch t-tests within and pooled across combinations. Yield per unit peak LAI was each plot’s yield divided by its maximum within-season LAI, from a locally weighted (LOWESS) fit to its LAI time course. The pooled yield comparison was reproduced from Tamang et al. (2026) for the same panel and not retested here. The narrow-minus-broad difference in canopy A (ΔA) was modeled as a quadratic function of the mean canopy LAI of narrow and broad plots, fitted by ordinary least squares to the 40 combination-by-date mean differences and summarized by R² and root-mean-square error (RMSE). Three alternative forms (linear, logistic and a three-parameter exponential rise to a plateau) were ranked alongside the quadratic by the Akaike information criterion (AIC; Fig S5). Field-validation contrasts used two-sided Welch t-tests by leaflet type on the per-plot light extinction coefficient (Beer k; Methods S1.9), total leaf area index, and leaf-area median height (LITES and SunScan) and, for LiDAR, on plot means of median foliage height, the share of returns above 70% of canopy height and canopy height over the canopy-closure window (DAP 70-75; 8 and 13 August 2024).

### Genotyping, heritability, and genome-wide association

Genotyping used the AgriPlex 1K SNP panel on 189 BC3 lines (1,327 SNPs; Tamang et al., 2026). The genotype matrix used here contained 1,253 of these markers. After quality control (call rate ≥ 0.80, minor allele frequency ≥ 0.05, and at least two genotype classes), 170 markers were retained and coded as {0, 1, 2} alternative-allele dosages. A genomic relationship matrix was constructed from the marker dosages (details in Methods S1). After overlap of the trait BLUPs (best linear unbiased predictors) with the kinship matrix, 188 lines were retained for all four traits.

BLUPs per line were computed for four traits: leaflet length-to-maximum-width ratio (Leaflet L:W; model in Methods S1.7), yield divided by maximum LAI (Yield per LAI), reproductive-stage Helios-predicted canopy net photosynthesis (Canopy A), and reproductive-stage Helios-predicted canopy light-use efficiency (Canopy LUE). Reproductive-stage values were averaged over 45 < DAP ≤ 85 and LAI ≥ 3 (per-line BLUPs in Table S17). Yield per LAI, Canopy A and Canopy LUE used y = µ + Location + Spacing + (1 | Line) in *lme4* (Bates et al., 2015; R 4.5.2, R Core Team, 2025). Cullis broad-sense heritability was computed for each trait (Cullis et al., 2006; per-trait heritability in Table S18).

Marker-trait associations were tested with a kinship-corrected mixed model for the slope (rrBLUP::mixed.solve; Endelman, 2011) and ordinary least squares for the standard error and two-sided P-value. Significance was set at the Bonferroni threshold (P < 0.05/170). At the lead chromosome-20 locus (SNP1293, Gm20:34,667,185, within *GmJAG1*), per-allele effects were standardized as β_std = β/SD(BLUP) with 95% confidence intervals.

## Results

### Ground-truth phenotypes confirmed the near-isogenic contrast

We characterized the BC_3_F_2:4_ broad-and narrow-leaflet NIL panels at two sites, with per-plot canopy LAI, leaflet morphometrics, leaf gas exchange, and end-of-season grain yield measured as ground truth for modeling (Fig 1). Narrow-leaflet lines reached a lower peak canopy LAI than broad-leaflet lines (9.0 versus 10.2 m² m^-^², about 12% lower; Fig 1C). Their leaflets were longer (11.7 versus 10.1 cm), about 38% narrower (maximum width 4.9 versus 7.8 cm) and 29% smaller (41.0 versus 57.4 cm²), giving a mean length-to-maximum-width ratio of 2.4 versus 1.3 (Fig 1D; leaflet area in Fig S2D). Jmax25 was 5% higher in narrow than in broad lines (215 versus 205 µmol m^-^² s^-^¹; Welch t, P = 0.015; n = 164 plots), whereas Vcmax25 did not differ significantly (146 versus 142 µmol m^-^² s^-^¹; P = 0.16; n = 198 plots) (Figs S1B, C).

### Narrow-leaflet canopies fixed more carbon

Three-dimensional ray-traced canopy A predictions showed narrow-leaflet canopies fixed more carbon than broad-leaflet canopies throughout the late vegetative and reproductive period at all four location x row spacing combinations (Fig 2A; illustrative canopy renderings in Fig 2B). At the first measurement date (mean LAI 0.7-1.6), narrow-leaflet canopies had slightly lower canopy A than broad-leaflet canopies in every combination (narrow-minus-broad −0.7 to −1.0 µmol CO_2_ m^-^² s^-^¹; Wilcoxon rank-sum P ≥ 0.18), when narrow plots also had about 10% less leaf area. From the second date onward, the narrow-leaflet advantage was positive at every date and grew as the canopy filled, peaking after canopy closure. Within each combination, peak canopy A was greater in narrow than in broad-leaflet lines (Wilcoxon rank-sum P ≤ 7.3 x 10^-^¹¹ in all four combinations), with narrow-minus-broad differences of +2.14 (SF 38 cm), +3.95 (SF 76 cm), +3.81 (EF 38 cm), and +2.90 µmol CO_2_ m^-^² s^-^¹ (EF 76 cm). Per-combination Cohen’s *d* ranged from −1.26 (EF 76 cm) to −3.68 (SF 76 cm).

**Figure 2.**
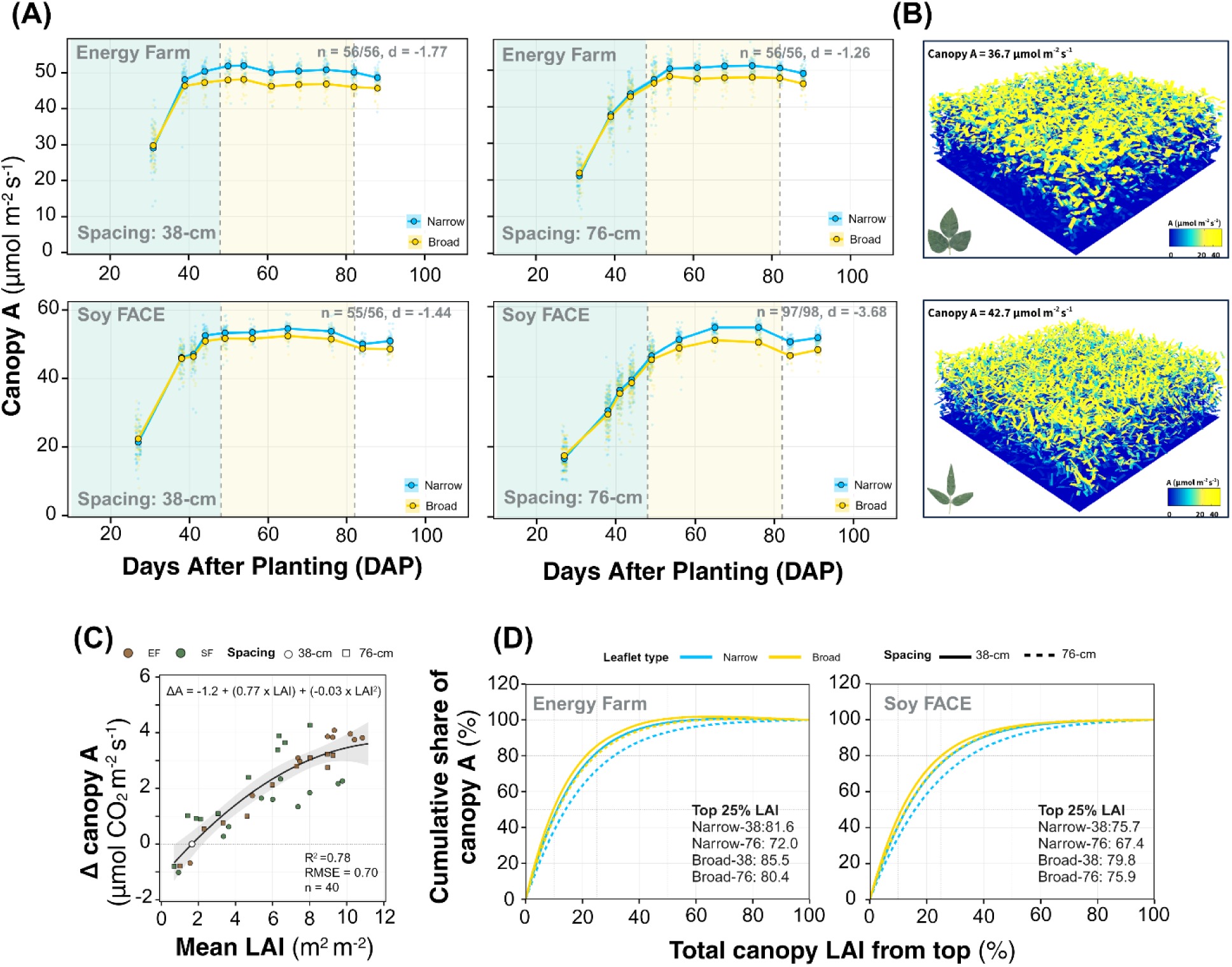
Helios-predicted canopy net photosynthesis (canopy A) of broad- and narrow-leaflet soybean near-isogenic lines at two sites and two row spacings. **(A)** Seasonal course of canopy A against days after planting (DAP) at the Energy Farm (top) and SoyFACE (bottom) in 38-cm (left) and 76-cm (right) rows. Small points are plots, lines and ribbons are means ± SE of plots, and shading marks growth stages as in Fig 1C. Each panel gives n (narrow/broad plots) and Cohen’s d (broad minus narrow, pooled SD) for per-plot peak canopy A. **(B)** Illustrative Helios canopies of broad (top) and narrow (bottom) leaflets, built from the mean leaf traits of the Energy Farm 76-cm plots on 11 August 2024 (DAP 73) with the same number of leaflets per square meter. Each sub-leaf element is colored by its net photosynthesis, and the canopy total is given above each canopy. **(C)** Narrow-minus-broad difference in canopy A (ΔA) against the mean leaf area index (LAI) of narrow and broad plots, with one point per site x row spacing x date (n = 40) and a quadratic fit with its 95% confidence band. The open circle marks ΔA = 0. **(D)** Cumulative share of canopy A against cumulative share of canopy LAI from the canopy top at the Energy Farm (left) and SoyFACE (right), pooled across the plots of each leaflet type and row spacing at their peak-LAI date. Inset values give the share of canopy A in the top 25% of LAI. Shares above 100% occur where leaves lower in the canopy had net respiration.

The advantage of narrow leaflets scaled with mean canopy LAI (Fig 2C). Modeling the difference as a quadratic function of mean canopy LAI gave ΔA = −1.19 + 0.77·LAI −0.031·LAI² (R² = 0.78, RMSE = 0.70 µmol CO_2_ m^-2^ s^-1^, n = 40 location x row spacing x DAP combinations), with ΔA = 0 at canopy LAI = 1.65 (1.0-2.5 when the four combinations were fitted separately). ΔA increased across the full observed LAI range and the fitted quadratic’s maximum (+3.66 µmol CO_2_ m^-^² s^-^¹ at LAI ≈ 12.5) lay beyond the highest observed mean LAI of 10.8. An exponential rise to a plateau fitted equally well (ΔAIC = 0.05) and crossed zero at LAI = 1.64, while the linear and logistic forms fitted less well (ΔAIC = 2.6 and 3.0; Fig S5).

The leaflet types also differed in where canopy A was fixed (Fig 2D). In broad-leaflet canopies, 76-85% of canopy A occurred within the top 25% of LAI, compared with 67-82% in narrow-leaflet canopies, a difference of about 4 percentage points in 38-cm rows and 8.5 in 76-cm rows. Net-respiring leaves at the bottom of the canopy reduced canopy A by 2% or less in all four combinations.

### Narrow-leaflet canopies used equal light more efficiently and spread it more evenly

Unlike canopy A, light absorption was similar in both leaflet types. After canopy closure (DAP ≥ 70), broad-and narrow-leaflet canopies absorbed 84-93% of PAR (per-plot means; Fig S6).

Narrow-minus-broad differences in post-closure mean fAPAR were +0.004 (SF 38 cm), −0.001 (SF 76 cm), +0.004 (EF 38 cm) and +0.003 (EF 76 cm), all within a ±0.02 equivalence margin (TOST, P < 0.001 in each combination; Fig S6B; seasonal trajectories in Figs 3A, S7A). Per-plot peak fAPAR differed by 0.005 or less between leaflet types (Wilcoxon rank-sum P = 0.042, 0.008, 0.039 and 0.76 for SF 38 cm, SF 76 cm, EF 38 cm and EF 76 cm; Cohen’s *d* from −0.37 to +0.47), small relative to the Cohen’s *d* observed for canopy A. Before canopy closure, narrow-leaflet canopies had slightly lower fAPAR than broad-leaflet canopies, by 0.03 at the first measurement date in every combination (Figs 3A, S7A).

**Figure 3.**
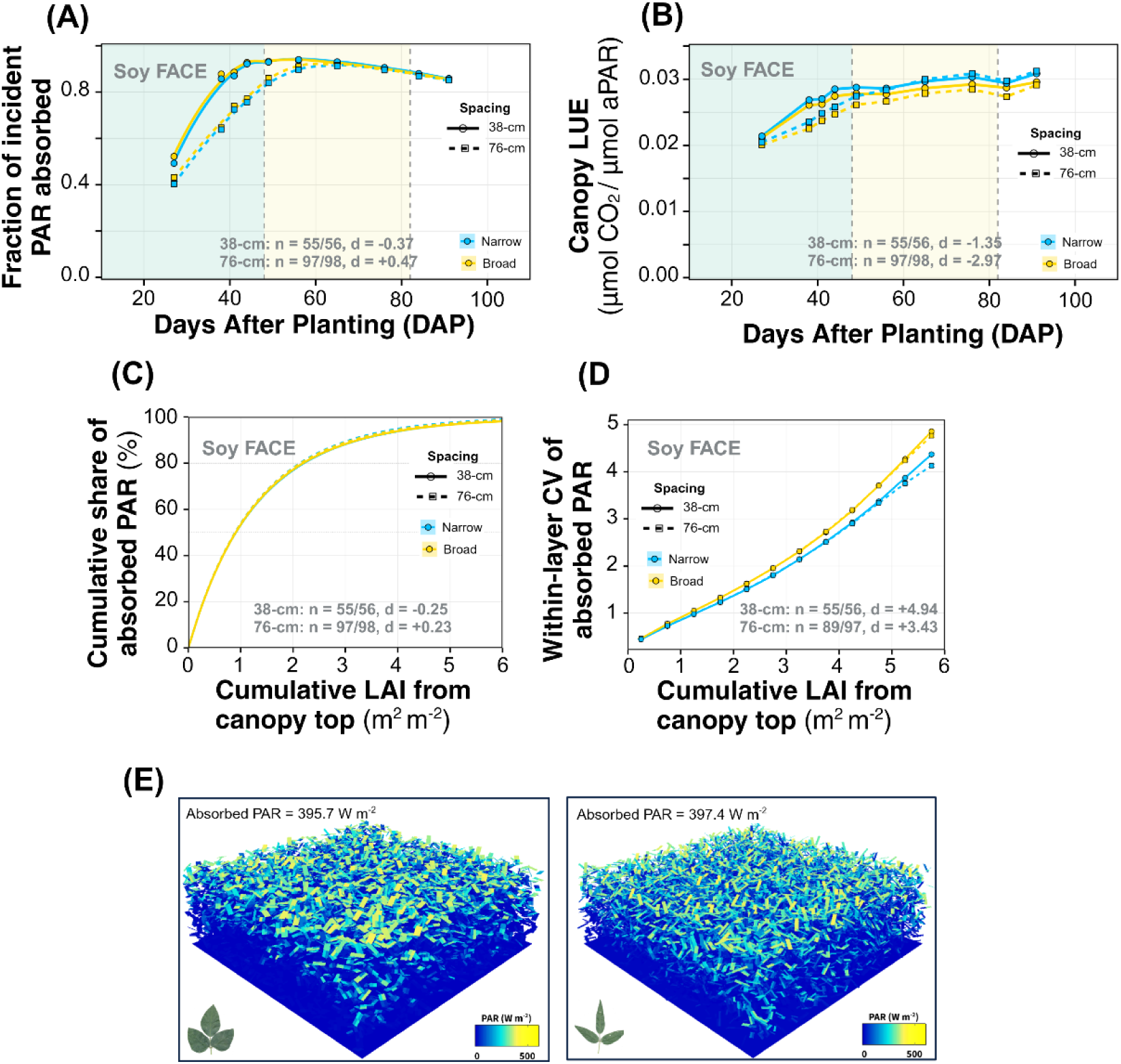
Helios-predicted light absorption, light-use efficiency and within-canopy light distribution in broad- and narrow-leaflet soybean near-isogenic lines. **(A)** Fraction of incident photosynthetically active radiation (PAR) absorbed (fAPAR) and **(B)** canopy light-use efficiency (LUE, canopy net photosynthesis per unit absorbed PAR) against days after planting (DAP) at SoyFACE, with growth-stage shading as in Fig 1C. **(C)** Cumulative share of each plot’s absorbed PAR against cumulative leaf area index (LAI) from the canopy top (means of per-plot curves). **(D)** Leaf-area-weighted coefficient of variation (CV) of absorbed PAR among sub-leaf elements in successive 0.5-LAI layers (mean ± SE of plots). C and D show SoyFACE on 19 August 2024 (DAP 76), the peak-LAI date of all four leaflet type x row spacing classes. **(E)** The illustrative canopies of Fig 2B (broad left, narrow right) colored by absorbed PAR, with each canopy’s total. Panels A-D give n (narrow/broad plots) and Cohen’s d (broad minus narrow, pooled SD) for per-plot peak fAPAR (A), peak LUE (B), the cumulative LAI holding half of the absorbed PAR (C) and the mean CV to a cumulative LAI of 6 (D, plots with LAI ≥ 6). Energy Farm counterparts of A-D are in Fig S7.

Despite absorbing similar fractions of incident PAR, narrow leaflet canopies converted absorbed light to carbon at higher rates than broad-leaflet canopies (Figs 3B, S7B). Peak canopy LUE was higher in narrow than broad-leaflet lines at both locations and row spacings (Wilcoxon rank-sum P < 10^-^¹⁰ in each), with per-combination Cohen’s *d* ranging from −1.30 to −2.97. The advantage corresponded to a 4 to 7% relative increase in canopy LUE (+0.0013 to +0.0021 µmol CO_2_ µmol^-^¹ absorbed PAR in absolute terms).

At the same cumulative LAI from the canopy top, absorbed PAR followed the same vertical profile in both leaflet types, with half of each plot’s absorbed PAR within a cumulative LAI of 0.90-0.94 (Wilcoxon rank-sum P ≥ 0.19 in each combination; Figs 3C, S7C). Within each depth, however, absorbed PAR was spread more evenly among leaves in narrow-leaflet canopies. The coefficient of variation of absorbed PAR among sub-leaf elements was lower than in broad-leaflet canopies in every 0.5-LAI layer down to a cumulative LAI of 6, in all four combinations with the gap widening with depth (mean over layers 2.11-2.16 vs 2.29-2.35; P < 10^-17^ in each; Figs 3D, S7D). Illustrative renderings of broad-and narrow-leaflet canopies colored by absorbed PAR are shown in Fig 3E.

### The advantage was rooted in three-dimensional canopy structure

To separate the architectural contribution from leaf physiology, we compared Helios with a one-dimensional sun-shade model parameterized with identical plot-level biochemistry, so that leaf-type differences in Vcmax25 and Jmax25 (Figs S1B, C) entered both models. The 1D and 3D models were strongly correlated across all plot and date combinations (Pearson r = +0.99, RMSE = 2.15 µmol CO_2_ m^-^² s^-^¹, n = 5,299 plot-date combinations; Fig 4A). Averaged over all measurement dates, Helios predicted narrow-minus-broad differences in canopy A of +1.31 to +3.04 µmol CO_2_ m^-^² s^-^¹ across the four location x row spacing combinations, whereas the 1D model predicted differences of −0.84 to +0.88 µmol CO_2_ m^-^² s^-^¹ (Fig 4B; Table S19). The architectural residual, the Helios difference minus the 1D difference, was positive in every combination (+1.75 to +2.17 µmol CO_2_ m^-^² s^-^¹). It equaled 71% (EF 38 cm) and 85% (SF 76 cm) of the Helios difference and exceeded it at SF 38 cm and EF 76 cm, where the 1D model predicted slightly lower canopy A for narrow than for broad lines. Using only the 164 plots with both Vcmax25 and Jmax25 measured, the residual ranged from +1.71 to +2.04 µmol CO_2_ m^-^² s^-^¹.

**Figure 4.**
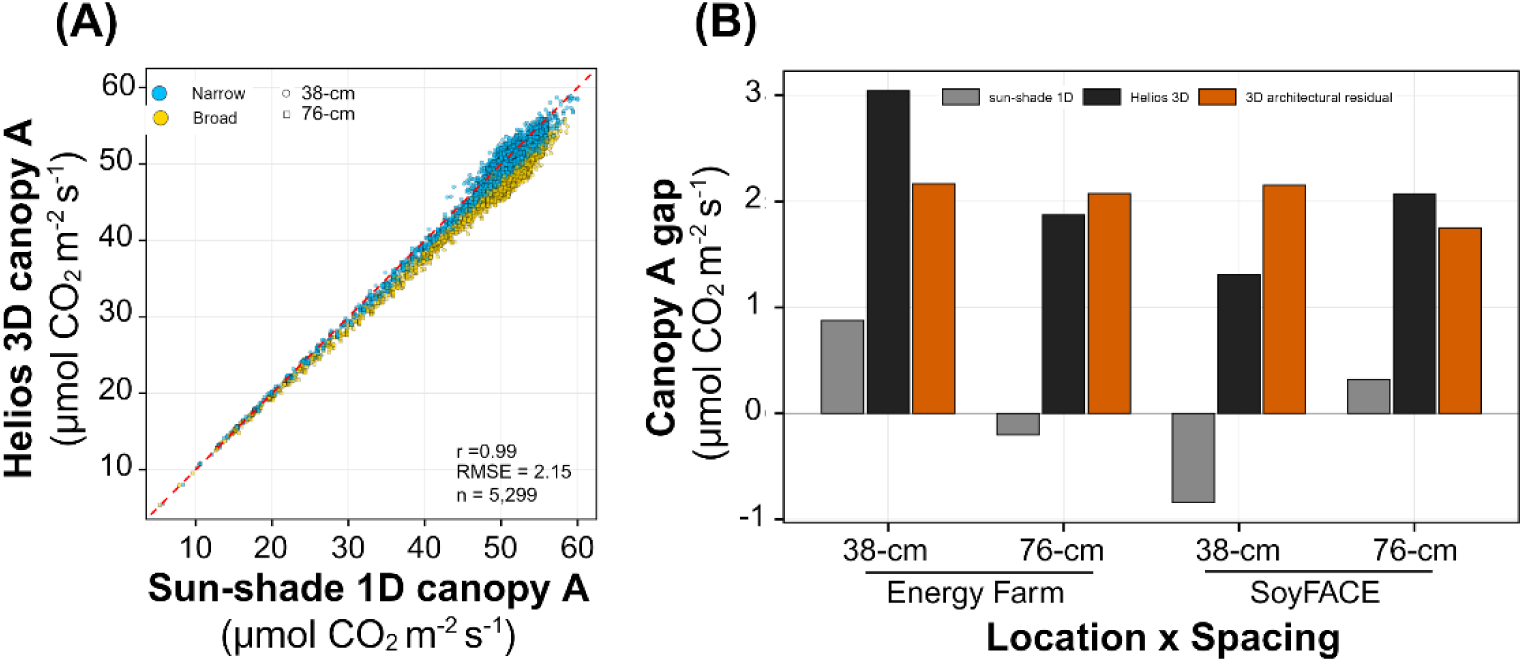
Three-dimensional (3D) Helios versus one-dimensional (1D) sun-shade predictions of canopy net photosynthesis (canopy A) in broad- and narrow-leaflet soybean near-isogenic lines. **(A)** Helios against 1D canopy A for all 5,299 plot-dates, with the 1:1 line dashed. The panel gives Pearson r and the root-mean-square error (RMSE). **(B)** Narrow-minus-broad difference in canopy A, averaged over all dates in each site x row spacing combination, predicted by the 1D model and by Helios, and their difference, Helios minus 1D (the 3D architectural residual). Values are in Table S19.

Leaflet length times average width, the Helios leaflet inputs, reproduced measured leaflet area (n = 530, R² = 0.997, slope = 0.996; Fig S8A). In a test of leaflet subdivision, going from the 9 elements per leaflet used in the main simulations to 225 elements lowered canopy A by 12-17% and the narrow-minus-broad difference by 25-52% (Fig S8B, C; Table S12), but narrow lines had higher canopy A than broad lines at every subdivision in all four location x row spacing combinations. Across plots, canopy A was not related to the number of leaflets in the simulated canopy (all slopes P > 0.08; R^2^ ≤ 0.051; Fig S8D).

### Field measurements matched the predicted light profile and showed small differences in leaf-area distribution

We compared the predicted within-canopy distributions with independent field measurements at both sites (Fig 5). At SF (76-cm rows, DAP 77-79), the Helios absorption profile and the LITES interception profile both rose with cumulative leaf area from the canopy top and leveled off toward the canopy floor (Fig 5A). Measured interception exceeded predicted absorption at the canopy floor by +0.092 (SD 0.018), and the difference was positive in 30 of 31 plots. Light extinction did not differ between narrow-and broad-leaflet lines (per-plot Beer k = 0.94 vs 0.89, P = 0.69). SunScan leaf-area profiles from the same plots showed that broad-leaflet canopies had more total leaf area than narrow-leaflet canopies (LAI 7.2 vs 6.1, P < 0.001) and that their leaf area was centered higher in the canopy (leaf-area median height 56% vs 52% of canopy height, P = 0.049; Fig 5B).

**Figure 5.**
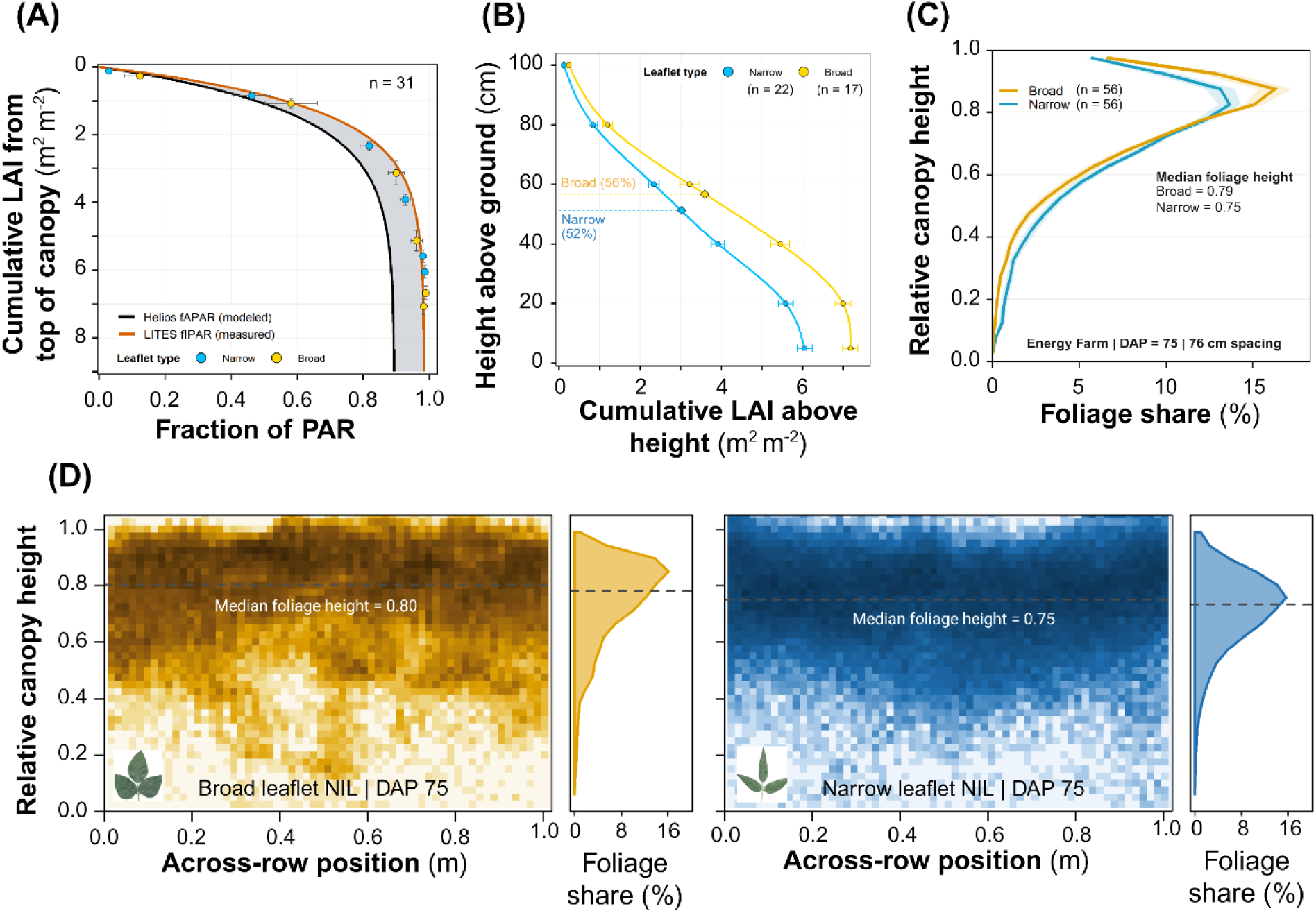
Predicted and measured within-canopy light and leaf-area distribution in broad- and narrow-leaflet soybean near-isogenic lines (NILs) at two sites. **(A)** Helios-modeled fraction of photosynthetically active radiation (PAR) absorbed (fAPAR) and the fraction of PAR intercepted (fIPAR), measured with pole-mounted quantum sensors (LITES), against SunScan cumulative leaf area index (LAI) from the canopy top, at SoyFACE in 76-cm rows 77-79 days after planting (DAP). The shaded band is canopy reflectance to sky, R_sky = fIPAR - fAPAR ∼ 0.09. Points are class means (± SE on both axes) at six heights from 33 NIL plots (22 Narrow, 11 Broad), and the Helios curve uses the 31 of these simulated on DAP 76. **(B)** Height against SunScan cumulative LAI above that height for 22 Narrow and 17 Broad plots, including six plots of the broad-leaflet recurrent parent LD11-2170 (class means ± SE, spline fits). Diamonds mark each class’s leaf-area median height. **(C)** Share of overhead LiDAR foliage returns by relative canopy height at the Energy Farm (76-cm rows) at canopy closure (DAP 75), as class means ± SE (n = 56 per class). **(D)** LiDAR return density across the row and with relative height for one broad- and one narrow-leaflet plot on DAP 75, with each plot’s height profile and median foliage height (dashed line).

Overhead LiDAR at EF provided an independent structural measurement at the second site (76-cm rows, 56 Broad and 56 Narrow NILs). At canopy closure (DAP 70-75), the median height of foliage relative to canopy height was greater in broad-leaflet than in narrow-leaflet canopies (0.80 vs 0.76, P = 0.019), and a larger share of LiDAR returns lay in the upper 30% of the canopy (0.74 vs 0.67, P = 0.020; Fig 5C). Canopy height did not differ (0.81 vs 0.77 m, P = 0.11; Table S9; representative plots in Fig 5D). Before canopy closure, median foliage height did not differ between leaflet types or was slightly greater in narrow-leaflet canopies (DAP 32-36; Table S8).

### Narrow lines produced equivalent yields with less leaf area

Grain yield was equivalent between broad and narrow-leaflet lines (5,801 versus 5,756 kg ha^-^¹, P = 0.43; Tamang et al., 2026). Per-plot mean differences in yield across the four location x row spacing combinations ranged from −129 to +112 kg ha^-^¹ with no consistent direction (Welch t per combination, P ≥ 0.083). Per-plot peak leaf area index was 7-15% lower in narrow than in broad lines in every combination (Welch t, P < 0.0001; Fig 1C).

Because narrow lines produced equivalent yield from less leaf area, their grain yield per unit leaf area was higher than that of broad lines (Fig 6A, B). The narrow advantage in yield per unit peak leaf area was significant in every location x row spacing combination (Welch t test on per-plot ratios, P ≤ 0.0034 in all four combinations; per-combination Cohen’s *d* ranged from −0.57 to −1.37). Pooled across all 515 plots with paired yield and peak leaf area records, the narrow advantage in yield per unit peak leaf area was +80.4 kg ha^-^¹ per LAI (+13.7%, Welch t P < 0.0001, pooled Cohen’s *d* = −0.96).

**Figure 6.**
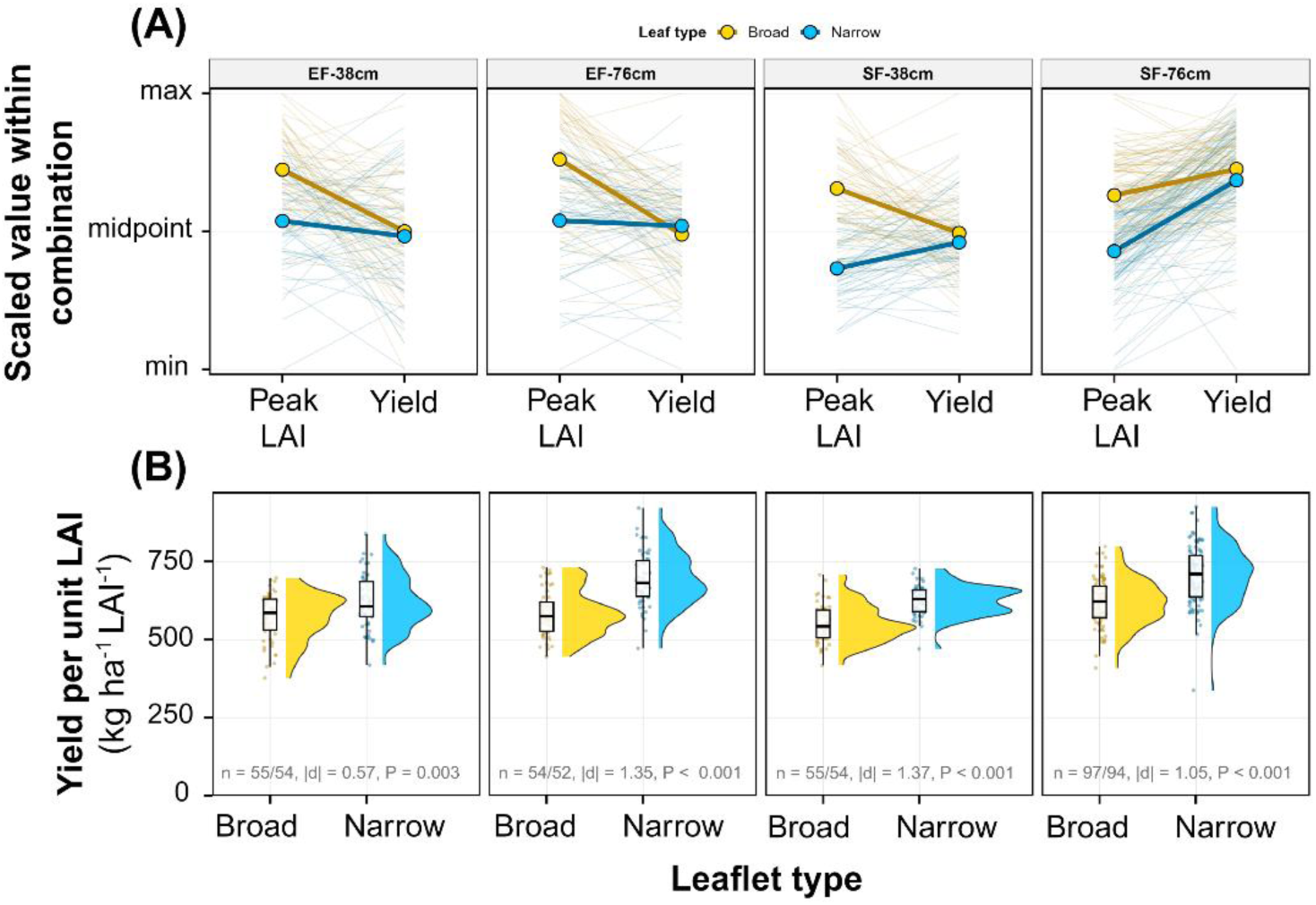
Peak leaf area index (LAI) and grain yield of broad- and narrow-leaflet soybean near-isogenic lines in each site x row spacing combination (n = 515 plots). **(A)** Each plot’s peak within-season LAI and grain yield, scaled from 0 (minimum) to 1 (maximum) within the combination and joined by a thin line. Thick lines join the leaflet-type means. **(B)** Per-plot grain yield per unit peak LAI by leaflet type, shown as density, box plot (median and interquartile range) and points. Each panel gives n (broad/narrow plots), Cohen’s |d| and the Welch t-test P.

### Leaflet shape, canopy net photosynthesis, canopy LUE and yield-per-leaf-area mapped to *GmJAG1*

For the four traits (line BLUPs in Fig 7A), Cullis broad-sense heritability was 0.95 for Leaflet L:W, 0.56 for Canopy A, 0.62 for Canopy LUE, and 0.39 for Yield per LAI (per-trait heritability in Table S18).

**Figure 7.**
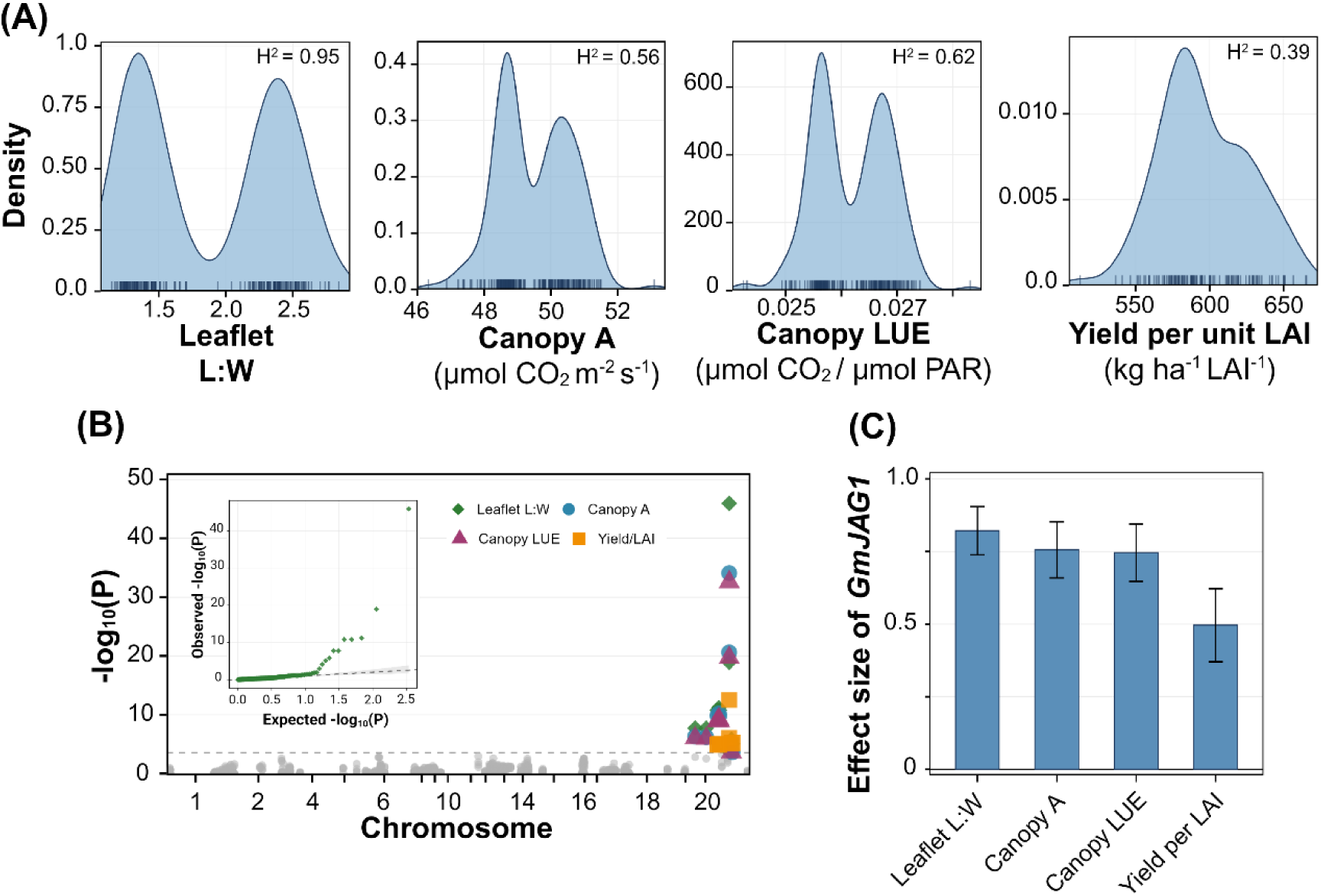
Genome-wide association analysis of leaflet shape, Helios-predicted canopy traits and yield per leaf area in 188 BC₃F₂:₄ near-isogenic lines. **(A)** Distributions of line best linear unbiased predictors (BLUPs) for leaflet length-to-maximum-width ratio (Leaflet L:W), reproductive-stage canopy net photosynthesis (Canopy A), canopy light-use efficiency (Canopy LUE) and yield per unit peak leaf area index (Yield per LAI), with each trait’s broad-sense heritability (H²). **(B)** Manhattan plot of −log₁₀(P) along the genome for the four traits. Markers above the Bonferroni threshold (dashed line, P = 0.05/170) are colored by trait, and the inset is the quantile-quantile plot for Leaflet L:W. **(C)** Standardized per-allele effect (β divided by the SD of the BLUPs) at SNP1293 (Gm20:34,667,185, within GmJAG1) for each trait, with 95% confidence intervals.

Genome-wide association on BLUPs identified a single locus on chromosome 20, SNP1293 at Gm20:34,667,185 within *GmJAG1*, as the top marker for all four traits (Fig 7B). Top-marker P-values were 1.16 x 10^-^⁴⁶ for Leaflet L:W, 8.14 x 10^-^³⁵ for Canopy A, 2.67 x 10^-^³³ for Canopy LUE, and 3.13 x 10^-^¹³ for Yield per LAI; each was below the Bonferroni threshold of 2.94 x 10^-^⁴ (top marker per trait in Table S20).

The standardized per-allele regression coefficient at SNP1293 clustered near 0.75-0.85 SD per allele for Leaflet L:W (β_std = +0.82, 95% CI [+0.74, +0.90]), Canopy A (β_std = +0.76, [+0.66, +0.85]), and Canopy LUE (β_std = +0.75, [+0.65, +0.84]), and was smaller for Yield per LAI (β_std = +0.50, [+0.37, +0.62]) (Fig 7C). All four 95% confidence intervals excluded zero.

## Discussion

Canopy carbon gain in field-grown crops is determined jointly by leaf-level photosynthetic capacity and by the spatial distribution of light within the canopy (Yin & Struik, 2015; Slattery & Ort, 2021). Of the two, the leaf photosynthetic capacity has received intensive attention from genetic-engineering programs (Croce et al., 2024; Long et al., 2025). To test the role of light distribution, we used the narrow-leaflet trait of soybean, a single-gene change in canopy architecture. Using three-dimensional ray-traced canopy biophysics simulations parameterized with per-plot field measurements from NILs at two locations and two row spacings, we addressed four questions. We asked whether the leaflet-shape effect on canopy carbon gain operates through light interception or canopy light-use efficiency and where in the canopy the difference originates. We then asked what fraction of the architectural advantage requires three-dimensional resolution to detect, and whether variation in the predicted canopy traits traces to the locus (*GmJAG1*) controlling the input morphology. Because our conclusions rest on model predictions, we anchored them with independent field measurements at both sites, which matched the predicted light profile and showed small differences in leaf-area distribution.

### Leaflet shape modulates canopy light-use efficiency

After canopy closure, both leaflet types absorbed similar fractions of incident PAR, so the higher carbon gain of narrow canopies came from converting that light 4 to 7% more efficiently (Fig 3B). This agrees with the greater canopy light-and nitrogen-use efficiency reported by Zhao et al. (2023) for lanceolate versus ovate cultivars. It also fits the higher intercepted PAR per unit leaf area reported by Bianchi et al. (2020) for lanceolate versus ovate NILs. Within the canopy, broad canopies fixed a larger share of their carbon in the upper 25% of canopy LAI (Fig 2D), mainly because they carry more leaf area. Absorbed PAR declined with cumulative LAI from the canopy top in the same way in both leaflet types (Fig 3C), so the upper 25% of a broad canopy’s LAI spans more of the well-lit canopy. The leaflet types differed instead in how absorbed PAR was spread among leaves at the same depth. Narrow canopies spread it more evenly in every layer (Fig 3D), keeping more leaves in the efficient linear region of the A-PAR response.

This within-layer difference, set by the smaller projected area of individual narrow leaflets, is resolved only in three dimensions. Because leaf photosynthesis saturates with light, canopy carbon gain depends on how evenly absorbed PAR is spread among leaves as well as on how much is absorbed (de Pury & Farquhar, 1997; Long et al., 2006). A modest leaf-biochemistry contribution is also present, as narrow-leaflet plots had higher Jmax (Fig S1C), consistent with the greater area-based photosynthetic capacity of lanceolate leaves (Tamang et al., 2023).

In the field, broad canopies held their foliage slightly higher by about 4% of canopy height (Fig 5B, C). This difference is small, so placing leaves at random through the canopy height in the simulations is a reasonable approximation for both leaflet types. Narrow canopies carry less leaf area and hold it lower, so less foliage sits above their mid-canopy, and more light should reach it. This fits the higher red-to-far-red ratio that Bianchi et al. (2020) measured in the middle of lanceolate canopies.

### Detecting the architectural advantage required three-dimensional resolution

The narrow-versus-broad canopy A advantage crossed from slightly negative to positive at a canopy LAI of approximately 1.7 and increased with greater LAI (Fig 2C). Once mutual shading dominates within-canopy light distribution, the partitioning of absorbed PAR among leaves becomes a consequential canopy trait (Long et al., 2006). In the field, the LiDAR-measured difference in foliage distribution between leaflet types appeared only after canopy closure (Fig 5C; Table S8), when the simulated advantage was largest.

Unlike Helios, a one-dimensional model recovered little or none of the narrow advantage at any combination, so the architectural residual accounted for most or all of the advantage of narrow-leaflet canopies (Fig 4B). The one-dimensional model accounts for leaflet shape only through total LAI, applying the same vertical extinction to both leaflet types and collapsing all within-layer horizontal structure. The residual is therefore consistent with the more even spread of absorbed PAR among leaves at each depth in narrow canopies (Fig 3D). Other lower-dimensional canopy models share this limitation (Gu et al., 2022), as do crop growth models that use a single, constant extinction coefficient, the most common approach in wheat models (Liu et al., 2021). These models would miss the benefit of narrow leaflets.

### A canopy mechanism for yield parity in narrow-leaflet lines

Earlier work showed that lanceolate canopies operate at higher efficiency than ovate canopies (Bianchi et al., 2020; Zhao et al., 2023; Tamang et al., 2026). Our simulations offer a mechanistic explanation for the yield parity of narrow-and broad-leaflet lines (Mandl & Buss, 1981; Wells et al., 1993; Dinkins et al., 2002) despite the lower canopy leaf area of narrow lines (Fig 1C; Tamang et al., 2026). After canopy closure, narrow canopies absorb a similar fraction of PAR from less leaf area because, in both leaflet types, absorption is concentrated near the canopy top (half within the top unit of LAI; Fig 3C). They are also predicted to convert that light to carbon more efficiently, by spreading it more evenly among leaves (Fig 3D). This predicted carbon gain did not translate into higher yield, consistent with the midday clear-sky estimates being upper bounds on the season-integrated advantage (see Assumptions and limitations).

Other routes may also contribute to yield parity. Bianchi et al. (2020) proposed a photomorphogenic pathway, linking the higher mid-canopy red-to-far-red ratio of lanceolate canopies to more pod initiation. Our simulations did not resolve light quality, so this pathway may act alongside the efficiency gain. The narrow-leaflet allele also increases the number of seeds per pod (Jeong et al., 2012; Sayama et al., 2017). In this panel, pod numbers did not differ and broad lines had 9% heavier seeds (Tamang et al., 2026), so more seeds compensated for lighter seeds.

### Reduced canopy leaf area lowers metabolic investment without lowering yield

Sunlight is freely available in outdoor agricultural environments, but the canopy that intercepts it is built and maintained at metabolic cost. Each unit of leaf area is a recurring investment of fixed carbon, nitrogen, and water in construction and transpiration. A large share of leaf nitrogen in C3 leaves is invested in Rubisco, so radiation use efficiency depends on the balance between leaf-area expansion and leaf nitrogen content per unit area (Sinclair & Horie, 1989; Wright et al., 2004). In soybean, a field manipulation study showed that canopies carry more leaf area than is optimal. Removing a fraction of emerging leaves increased seed yield by 8% under both current and elevated [CO₂] (Srinivasan et al., 2017). Rising atmospheric CO₂ increases canopy leaf area further (Ainsworth et al., 2002; Dermody et al., 2006), which would widen this excess.

The narrow-leaflet trait reaches a similar end point genetically, producing equivalent grain yield from a lower peak LAI, a 13.7% advantage in yield per unit canopy leaf area. The additional leaf area of broad canopies extends the canopy into deeper shade, where little light remains to be absorbed (Fig 3C). Breeding has also reduced leaf size in soybean. Across Upper Midwest cultivars released over eight decades, mean leaflet area fell by 17% as yield rose, with no change in canopy coverage (Sreekanta et al., 2026). Because leaf area is the shared substrate for the canopy carbon, nitrogen, and water budgets, equivalent yield from less leaf area in narrow lines implies more economical use of those resources, although we did not measure canopy nitrogen or water use directly. The value of delivering equivalent grain from a smaller vegetative investment rises as nitrogen, water, and crop development time become more constrained by environmental conditions (Slattery & Ort, 2021).

### The canopy efficiency cascade traces to the *GmJAG1* leaflet-shape locus

Leaflet shape, canopy photosynthesis, canopy LUE, and yield per LAI all mapped to a single locus on chromosome 20, SNP1293, the marker for *GmJAG1* (Jeong et al., 2012; Fang et al., 2013; Cai et al., 2021). Because the near-isogenic lines differ mainly in the introgressed chromosome-20 segment, the significant markers span much of that chromosome (6.8-37.2 Mb; Fig 7B), so the scan localizes the introgression and cannot resolve the causal variant within it.

A second caveat applies to the simulated traits. Because canopy A and canopy LUE are biophysical functions of inputs (leaflet morphology and leaf biochemistry) that segregate at *GmJAG1*, their co-localization at the same locus is expected by construction and is not independent genetic evidence for the mechanism. The per-allele effect on Helios-predicted canopy physiology is nearly as large as the effect on directly measured leaflet shape (Fig 7C), so the simulation carries the leaflet-shape signal through to the canopy traits largely undiluted. By contrast, yield per LAI, measured in the field and independent of the simulation, had a smaller effect, in line with its lower heritability (Table S18). Its association with the locus reflects equal yield from lower LAI in narrow lines.

### Leaflet shape as a heritable architectural lever for canopy resource-use efficiency

Conversion efficiency is the principal remaining target for raising yield potential in major grain crops (Long et al., 2006; Zhu et al., 2010). That target has been pursued largely through modifying leaf biochemistry (Croce et al., 2024; Long et al., 2025). Our findings position canopy architecture as a complementary route for improving canopy conversion efficiency. This fits with global-scale evidence that canopy clumping architecture is a determinant of yield and nitrous oxide emissions across the major staple crops (Yan et al., 2026).

The narrow-leaflet trait suits this route well. It is heritable, controlled by a single locus (*GmJAG1*; Jeong et al., 2012; Fang et al., 2013), and has already been modified by gene editing (Cai et al., 2021). Because this single architectural change is predicted to improve canopy LUE without yield penalty in the BC_3_F_2:4_ background, it could be stacked with leaf-biochemistry improvements in breeding. Multi-environment confirmation across additional cultivar backgrounds and growing seasons will test whether the within-canopy redistribution mechanism generalizes across the soybean mega-environments in North America (Krause et al., 2025).

### Assumptions and limitations

Canopy A was evaluated at midday (12:00 CDT) under clear-sky direct and diffuse fluxes on each date, so the predicted narrow advantage reflects direct-beam conditions. Because diffuse radiation penetrates deeper and more uniformly than direct radiation (Li et al., 2014), the within-canopy redistribution underlying the advantage may weaken under overcast or highly diffuse conditions. This is consistent with Zhao et al. (2023), who attributed the lanceolate canopy advantage predominantly to the direct-light component. The midday clear-sky values therefore likely represent an upper bound on the diurnally and seasonally integrated advantage of narrow-leaflet canopies.

The simulations also assigned both leaflet types the same planophile leaf-angle distribution, so the architectural residual arises from leaflet dimensions alone. Because narrow leaflets were also smaller than broad leaflets (Fig S2D), the simulations do not separate the effect of leaflet elongation from that of leaflet size. Leaf-angle distribution was not measured. The field extinction comparison (Fig 5A) cannot reveal it, because the SunScan infers leaf area from transmitted light with an assumed leaf-angle distribution. If systematic differences in inclination were present, part of the effect attributed to leaflet shape could instead reflect shape-correlated differences in leaf angle.

In addition, the simulated canopies were horizontally homogeneous, without rows, so differences between 38-cm and 76-cm row spacings were represented only through measured LAI and plant height. The size of the predicted advantage depended on how finely leaflets were subdivided for ray tracing. Finer subdivision shrank the advantage without reversing it. At 225 instead of 9 elements per leaflet, the relative LUE advantage was 7-44% smaller (Fig S8B, C; Table S12), so the absolute advantages reported here are likely upper estimates.

Finally, the efficiency result itself is a model prediction. The field measurements could not resolve how absorbed PAR was spread among leaves at the same depth, the property associated with the simulated advantage. Direct canopy-scale gas-exchange measurements and type-specific leaf-angle distributions would confirm the predicted LUE advantage and refine these attributions.

## Author contributions

BGT designed the study, conducted the field experiments and measurements, ran the simulations, analyzed the data and wrote the manuscript. AD ran the LITES light-profile measurements. CJB led the SpiderCam LiDAR data collection. EAA supervised the project and revised the manuscript. All authors reviewed and approved the final manuscript.

## Acknowledgements

We thank Brett Feddersen for operating the SpiderCam LiDAR platform, Christopher Montes for help with planting, Jeanaflor Crystal T. Concepcion and Samuel Cheung for help with field data collection, Gregory Bernard for acquiring drone imagery, and Brian W. Diers and Sarah J. Schultz for developing the near-isogenic lines. Simulations were run on the Biocluster of the Carl R. Woese Institute for Genomic Biology, University of Illinois Urbana-Champaign.

## Funding

This work was supported by the project Realizing Increased Photosynthetic Efficiency (RIPE), funded by Gates Agricultural Innovations grant investment 57248, awarded to the University of Illinois, United States.

## Competing interests

None declared.

## Data availability

Helios source code is available at https://github.com/PlantSimulationLab/Helios under the GPLv3 license; simulations were run with commit c710cdfb (release v1.3.71). Per-plot input data, simulation outputs, the 1D sun-shade model implementation, and analysis scripts are deposited in Illinois Data Bank (https://doi.org/10.13012/B2IDB-9478251_V1). Analyses and figures used R 4.5.2 (R Core Team, 2025) and Python 3 (pandas, numpy, scipy, matplotlib).

## METHODS S1 - Supporting Information

### S1.1 A-Ci leaf sampling and handling

An undamaged, unshaded, recently matured trifoliate leaf (typically the fourth node from the top) from the middle rows of each plot was tagged the day before measurement, cut at the petiole base predawn, immediately submerged in water, and held in a cool, dark cabinet to maintain turgor. Before measurement, leaves were exposed to low light for at least 30 min. Leaf-type differences in the per-plot A-Ci estimates of Vcmax25 and Jmax25 were tested with two-sided Welch t-tests.

### S1.2 Barometric pressure

Barometric pressure was used as reported by each station. At SoyFACE, this was the sea-level-reduced value, about 2.6 kPa above the absolute pressure at the site. Pressure enters the simulations only through the clear-sky radiation model, in which this difference lowers incident PAR at SoyFACE by about 0.5%, equally for both leaflet types.

### S1.3 PROSPECT-D inversion

The per-plot parameters were leaf chlorophyll content (LCC, equivalent to PROSPECT-D Cab; µg cm^-^²) and leaf mass per area (LMA, equivalent to Cm; g cm^-^²); plots lacking direct SPAD or LMA received combination-mean substitution. The combination-level parameters (leaf structure N, carotenoids Car, equivalent water thickness Cw, anthocyanins Cant, and the senescent-pigment parameter Cs retained by the prosail Python implementation) were fitted separately for each of the four (location x row spacing) combinations by nonlinear least-squares inversion of that combination’s mean reflectance spectrum over 400-2400 nm, yielding four combination-specific parameter sets. LCC and LMA were held fixed at dataset-mean values during the combination-level inversion to avoid the known degeneracy of inverting all parameters jointly from reflectance alone (Jacquemoud & Baret, 1990; Féret et al., 2008, 2017). Each plot then received a forward run using its per-plot LCC and LMA together with the combination-specific N, Car, Cw, Cs, and Cant, returning R(λ) and T(λ) at 1-nm resolution from 400 to 2500 nm. The four scalar Helios inputs were obtained by arithmetic-mean band integration over 400-700 nm (PAR) and 700-2500 nm (NIR). Inversions and forward runs used the prosail Python package (version 2.0.5).

### S1.4 FvCB temperature dependencies and the coupled leaf solution

The Vcmax25 profile was computed for each sub-leaf element as l = (z -z_min)/(z_max -z_min) - 1, with z the element height. The coefficient 0.64 was taken from Wang et al. (2026). Temperature dependencies of Vcmax, the Michaelis-Menten constants of Rubisco for CO₂ and O₂, the CO₂ compensation point in the absence of mitochondrial respiration (Γ*), and dark respiration (Rd) followed Bernacchi et al. (2001); Jmax used the same form with activation energy corrected to 43.9 kJ mol^-^¹ (Sharkey et al., 2007). Atmospheric O₂ was 210 mmol mol^-^¹, PSII electron-transport quantum efficiency (α = 0.425 mol electrons mol^-1^ absorbed photons) and the light-response curvature (θ = 0.70) followed Bernacchi et al. (2003) and von Caemmerer (2000). Stomatal conductance used the simplified Buckley-Mott-Farquhar formulation (Buckley et al., 2003, 2012) with default Helios parameters, and boundary-layer conductance the Pohlhausen formulation. At each plot-DAP timestep the stomatal, energy-balance (leaf temperature and latent flux from absorbed shortwave + longwave radiation, sensible heat exchange, and transpiration), and radiation models were iterated in a two-pass solve: an initial pass at air-temperature leaf surfaces, then a second pass with leaf temperatures updated from the energy-balance solution. Per-leaf photosynthesis was computed once the system had converged. The eight combination-mean Rd25 values ranged 0.218-0.479 µmol CO₂ m^-^² s^-^¹ (Table S14); within-canopy variation in Rd25 was not represented.

### S1.5 Robustness of the per-parameter biochemistry substitution

The architectural-residual comparison was computed on the full set of 5,299 plot-DAP combinations. Restricting the comparison to the 164 plots with both parameters measured changed the per-combination architectural residual by at most 0.3 µmol CO₂ m^-^² s^-^¹ (full-set range +1.75 to +2.17; +1.71 to +2.04 when restricted to the 164 plots; Table S19), and the 3D-versus-1D correlation was unchanged (Pearson r = +0.99 in both cases). Per-(location x row spacing x leaflet shape) sample sizes are in Table S14.

### S1.6 One-dimensional model details

The 1D model used Spitters et al. (1986, Eq. 20) for direct/diffuse partitioning of incident PAR and the de Wit (1965) planophile leaf-angle distribution to match Helios. Two-stream beam-and-diffuse scattering followed de Pury & Farquhar (1997, Eqs. A2-A8), with the canopy beam-reflectance term (1 - ρ_cb) from Eq. A19. The canopy was discretized into ten depth layers with sun and shade leaf classes per layer; FvCB photosynthesis was applied to each class at each layer, and canopy net assimilation was the sum over layers of sunlit and shaded leaf assimilation, each multiplied by its leaf area in that layer. Leaf temperature and intercellular CO₂ were obtained from a two-pass iteration of the coupled stomatal-energy-balance solution using the same energy-balance and Buckley-Mott-Farquhar models, default coefficients, and Pohlhausen boundary-layer conductance as Helios; absorbed NIR driving the energy balance used the same two-stream scheme as PAR with each plot’s PROSPECT-D NIR optics. Settings that the 1D model requires but Helios does not define for a 1D canopy (leaf characteristic length, NIR share of shortwave, longwave exchange) were matched to Helios where possible. In sensitivity tests, other choices changed the architectural residual by ≤ 0.07 µmol CO₂ m^-^² s^-^¹. Predictions of the 1D model were aggregated to the (location x row spacing x leaflet shape) level for the Narrow-minus-Broad decomposition.

### S1.7 Genomic relationship matrix and BLUP extraction

Missing genotypes were mean-imputed within each marker for kinship-matrix computation only; per-marker association tests used non-imputed dosages. The genomic relationship matrix was K = WW′/p, where W is the marker-dosage matrix mean-centered within each marker and p the number of markers used. K was projected to the nearest positive-semidefinite matrix by clipping eigenvalues below 1 x 10^-^⁶ and reconstructing from the cleaned spectrum, then normalized by its mean diagonal. Per-line BLUPs were the line random effects, zero-centered for Leaflet L:W and with the model intercept added for the other three traits. For Leaflet L:W, per-plot length-to-maximum-width ratios were first spatially adjusted with a MAD Type II adjustment (Lin & Poushinsky, 1985), and per-line BLUPs were fitted in *lme4* to all plots, including checks, with y = µ + Location x Spacing + primary check (fixed) + (1 | Line) + (1 | Line:Environment), where Environment is the location x row spacing combination. Of the 204-line panel, 200 lines had computable BLUPs (Table S17), of which 188 overlapped the kinship matrix; the all-lines BLUPs for Yield per LAI, Canopy A, and Canopy LUE likewise comprised these 200 lines, retaining 188 after kinship overlap.

### S1.8 Plots used for the SoyFACE field validation

The LITES and SunScan analyses used 39 plots at SoyFACE: 22 narrow-leaflet NILs, 11 broad-leaflet NILs and six plots of the broad-leaflet recurrent parent LD11-2170. The LD11-2170 plots did not differ from the broad-leaflet NILs in Beer k, total LAI or leaf-area median height (Welch t, all P > 0.4), so they were pooled with the broad-leaflet NILs for comparisons between leaflet types (Table S21). With the broad-leaflet NILs alone, light extinction (P = 0.69) and total LAI (P = 0.002) gave the same results, and the leaf-area median-height difference was similar in size but not significant (55% vs 52% of canopy height, P = 0.18). Comparisons with Helios (Fig 5A) used only NIL plots, because LD11-2170 was not simulated.

### S1.9 Radiation-transport validation and canopy leaf-area structure

#### Radiation validation

For each LITES plot, the depth-resolved fraction of intercepted PAR, fIPAR(z) = 1 - PAR(z)/PAR(140 cm), was paired with the SunScan cumulative leaf area index above the same height at the six within-canopy heights (5-100 cm), so that the light and leaf-area axes came from separate instruments. The measured interception profile and the Helios-modeled absorption profile (cumulative fAPAR versus cumulative leaf area from the canopy top, for the 31 plots with a Helios simulation at DAP 76) were each summarized by a Mitscherlich fit, f = a(1 - exp(-b·L)), with L the cumulative LAI from the canopy top, by nonlinear least squares (LITES a = 0.984, b = 0.848; Helios a = 0.894, b = 0.761). Canopy reflectance to the sky was defined by the PAR mass balance 1 = fAPAR (absorbed) + R_sky (reflected) + T_floor (transmitted), so that R_sky(z) = fIPAR(z) - fAPAR(z) is the gap between the two profiles. Whole-canopy extinction by leaflet type was compared using per-plot Beer fits, fIPAR = 1 - exp(-k·LAI).

#### Leaf-area structure

Each plot’s SunScan cumulative-LAI-above profile was height-normalized using the manual plant height measured on 22 August 2024 (DAP 79). Two metrics were derived: total leaf area index (cumulative LAI above the canopy floor) and the leaf-area median height (the height above which half of the canopy leaf area lies), the latter expressed as a fraction of canopy height to separate vertical distribution from plant size.

### S1.10 Vertical and within-layer distribution of absorbed PAR

These measures used the Helios output for each sub-leaf element (height, area and absorbed PAR per unit leaf area) for every plot of each location x row spacing x leaflet type class, on the date of that class’s highest mean LAI, as in Fig 2D (dates in the Fig 3 and Fig S7 legends). Within each plot, elements were ordered by height from the canopy top, and cumulative LAI from the top was their summed area divided by the 9 m² simulated ground area. The vertical profile was summarized by L50, the cumulative LAI from the top at which summed absorbed PAR (absorbed PAR per unit leaf area x element area) first reached half of the plot’s total. L50 compares canopies at the same leaf area from the top, rather than at the same fraction of each canopy’s own leaf area as in Fig 2D. Per-plot curves of the cumulative fraction of absorbed PAR against cumulative LAI were evaluated on a 0.05-LAI grid and averaged across plots (Figs 3C, S7C). For the spread among elements at the same depth, each element was assigned to a 0.5-LAI layer (0-0.5, 0.5-1, …, 5.5-6) by its cumulative LAI from the top, and the coefficient of variation (CV) of absorbed PAR per unit leaf area among the elements in each layer was computed with element area as the weight (leaf-area-weighted standard deviation divided by leaf-area-weighted mean; Figs 3D, S7D). Each plot was summarized by the mean of the CVs of its twelve layers, for plots with total LAI ≥ 6. L50 and mean CV were compared between leaflet types within each location x row spacing combination by two-sided Wilcoxon rank-sum tests, with P from the normal approximation with continuity and tie corrections, and by Cohen’s *d* (pooled SD; positive = Broad > Narrow). Because CV increases as leaflets are divided more finely, CVs are comparable only at the same number of elements per leaflet. In the subdivision-sensitivity simulations (Table S12), element-level outputs at the same subdivision for both leaflet types (144 elements per leaflet) were available for Energy Farm 76 cm; there, the narrow-leaflet canopy had a lower CV than the broad-leaflet canopy in 11 of the 12 layers (mean 2.69 vs 2.93) and a similar L50 (0.93 vs 0.92).

### S1.11 LiDAR processing

The SpiderCam platform scanned at 450 mm s^-1^. Structural analysis used the data from 13 post-establishment flights (DAP ≥ 21). For each plot, point heights were divided by the plot’s 99th-percentile height to give relative canopy height. Foliage returns were those more than 0.05 m above the ground, and the metrics were computed on a standardized 1.5 m² plot footprint. Each LiDAR return is a point at which a laser pulse was reflected by a plant surface, so the share of returns at a given height shows where foliage is concentrated.

## Supplementary Tables

**Table S1.** Location, row spacing, leaflet type, line and field position of the 530 simulated plots.

**Table S2.** Plant height and leaf area index of each plot on each measurement date.

**Table S3**. Leaflet dimensions of each plot.

**Table S4**. Photosynthetic parameters from A-Ci curve fits for the 198 plots with curves.

**Table S5**. Meteorological inputs to the simulations for each site and measurement date.

**Table S6**. Grain yield and peak leaf area index of each plot.

**Table S7**. LITES light profiles and matched Helios absorption at SoyFACE (76-cm rows).

**Table S8**. LiDAR canopy structure of broad- and narrow-leaflet lines at the Energy Farm (76-cm rows), by flight date.

**Table S9**. LiDAR canopy-structure metrics of each plot at the Energy Farm (76-cm rows), by flight date.

**Table S10.** Leaf optical properties of each plot from PROSPECT-D.

**Table S11**. PROSPECT-D parameters fitted for each location and row spacing.

**Table S12**. Canopy A, fAPAR and canopy LUE at seven leaflet subdivisions (sub-leaf mesh test).

**Table S13.** Helios parameter inputs.

**Table S14**. Photosynthetic parameters by location, row spacing and leaflet type, as used in the simulations.

**Table S15**. Helios outputs and inputs for each plot and measurement date.

**Table S16**. One-dimensional sun-shade canopy A for each plot and measurement date.

**Table S17**. Line BLUPs for the four traits in the association analysis.

**Table S18**. Broad-sense heritability of the four BLUP traits.

**Table S19**. Helios and one-dimensional canopy A by location, row spacing, leaflet type and date, with the architectural residual.

**Table S20**. Association results at SNP1293 within GmJAG1 for the four traits.

**Table S21.** LITES and SunScan canopy-structure metrics of each plot at SoyFACE (76-cm rows).

---

**Figure S1.**
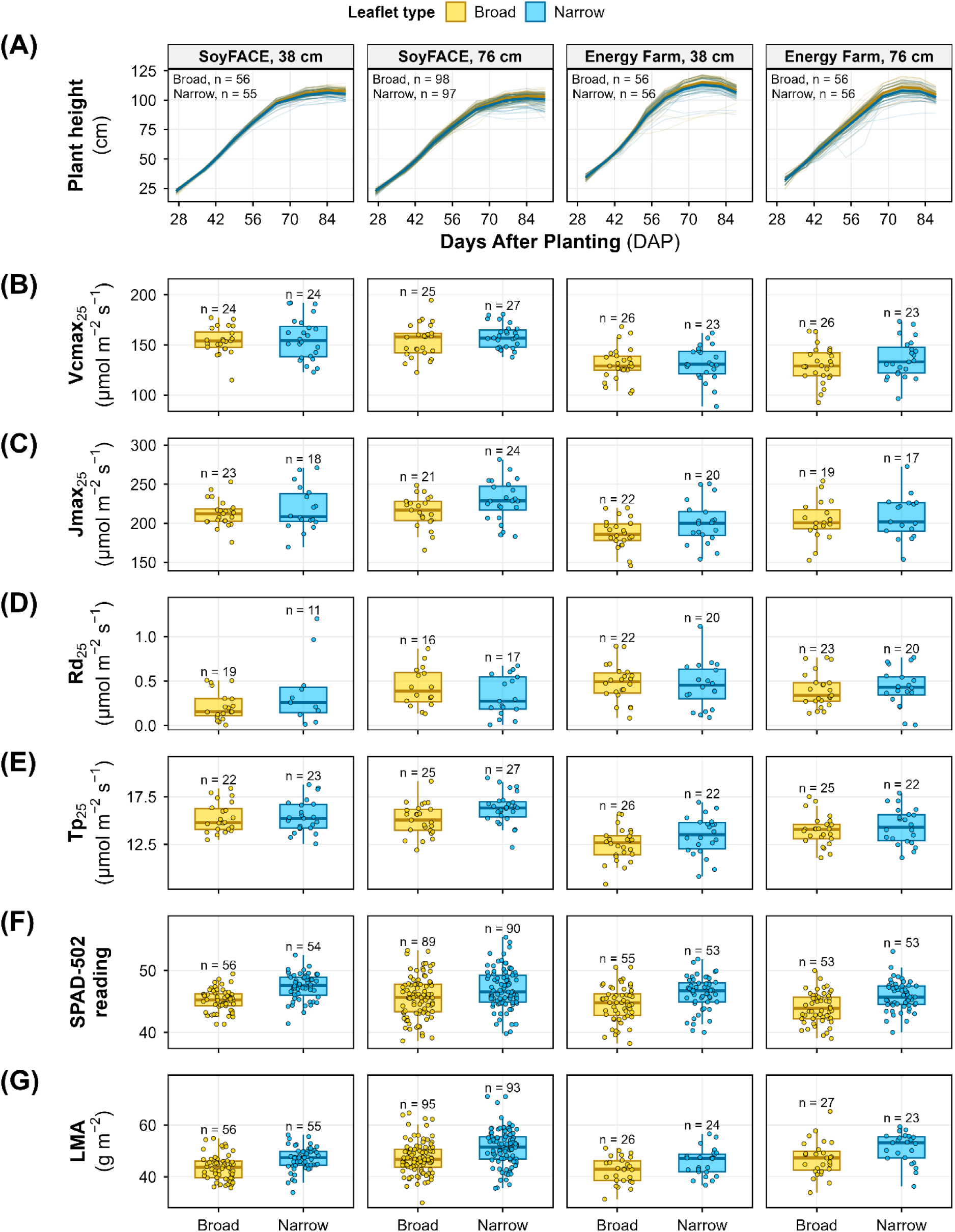
Per-plot inputs to the Helios simulations for broad- and narrow-leaflet soybean near-isogenic lines in each site x row spacing combination. **(A)** Plant height against days after planting (DAP), interpolated to each simulated date from field measurements. Thin lines are plots, thick lines and ribbons are leaflet-type means with 95% confidence intervals, and n is given in each panel. **(B)** Maximum carboxylation rate of Rubisco at 25 °C (Vcmax₂₅), **(C)** maximum electron transport rate at 25 °C (Jmax₂₅), **(D)** dark respiration at 25 °C (Rd₂₅) and **(E)** triose phosphate utilization at 25 °C (Tp₂₅), all from fits of leaf CO₂-response (A-Ci) curves. **(F)** SPAD-502 chlorophyll-meter readings. **(G)** Leaf mass per area (LMA), measured gravimetrically. Panels B-G show only plots with a direct measurement. Boxes show medians and interquartile ranges, and n is given above each box.

**Figure S2.**
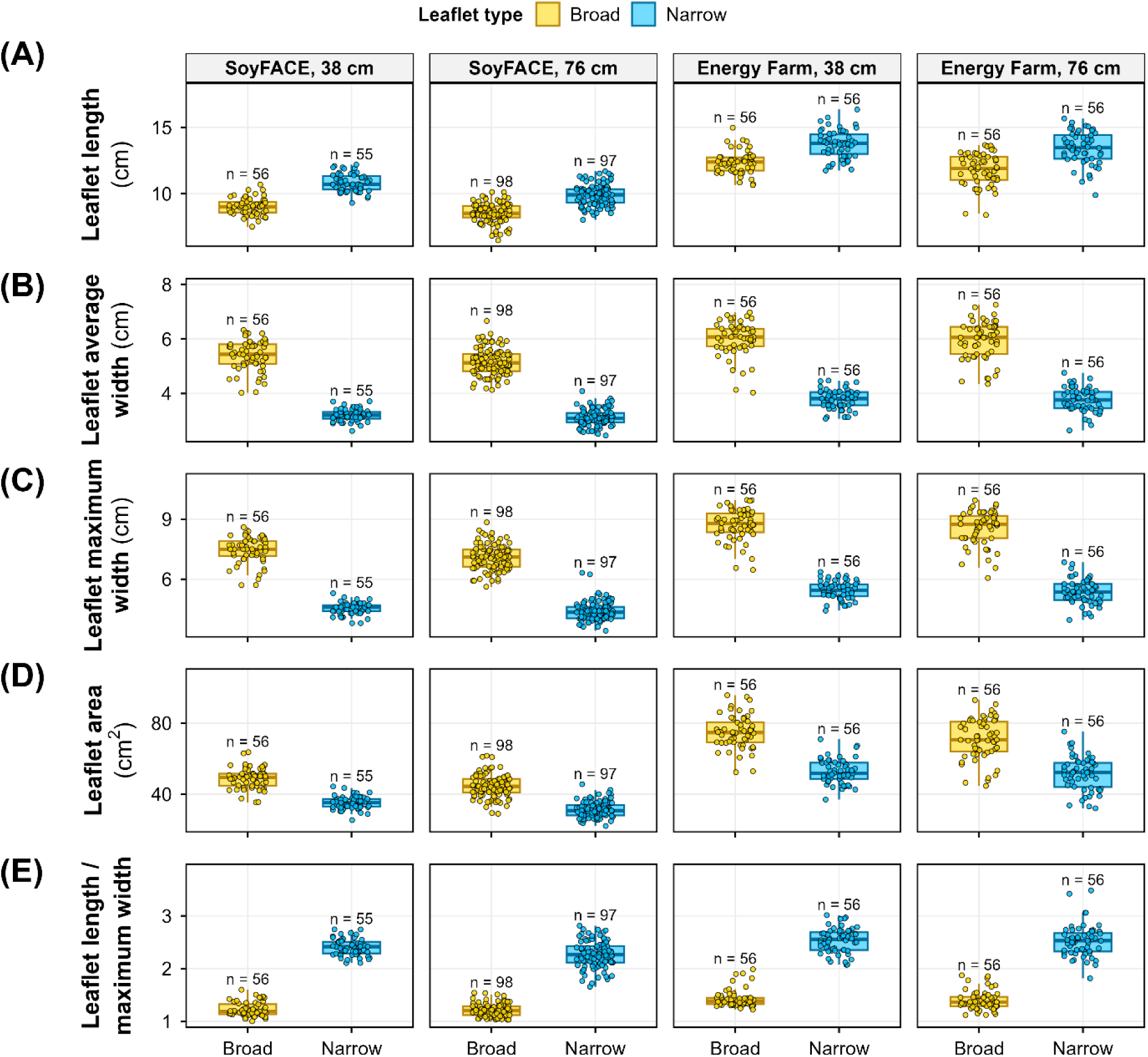
Per-plot leaflet dimensions of broad- and narrow-leaflet soybean near-isogenic lines in each site x row spacing combination. **(A)** Length, **(B)** average width, **(C)** maximum width, **(D)** area and **(E)** length-to-maximum-width ratio of leaflets, measured once per plot in 2024. Length and average width were the Helios inputs. Boxes show medians and interquartile ranges, and n is given above each box.

**Figure S3.**
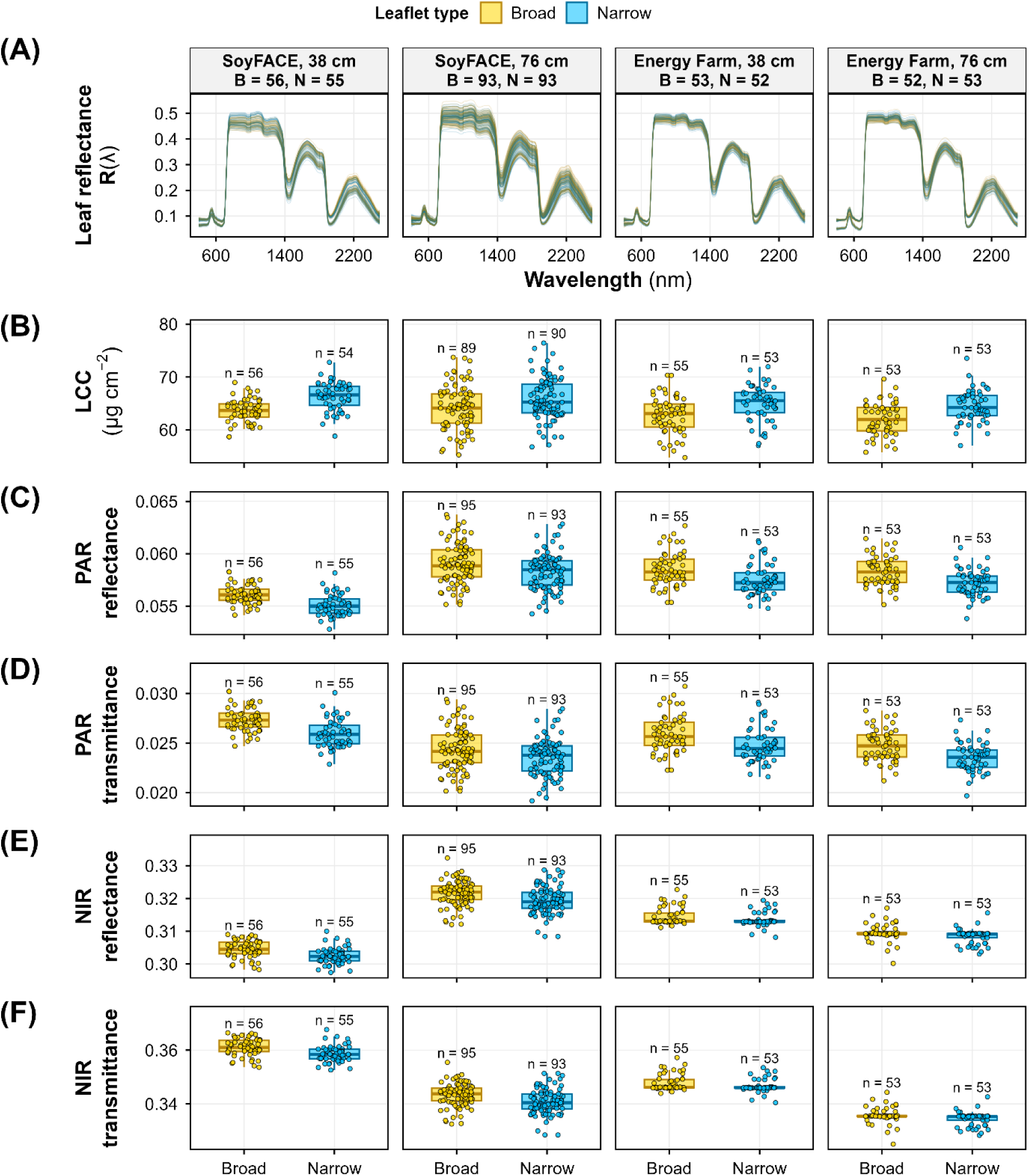
Leaf optical inputs to the Helios simulations for broad- and narrow-leaflet soybean near-isogenic lines in each site x row spacing combination. **(A)** Per-plot leaf reflectance spectra, R(λ), from 400 to 2500 nm, measured with a contact probe and averaged over the leaves measured in each plot. Panel headers give the number of broad (B) and narrow (N) plots. **(B)** Leaf chlorophyll content (LCC), from SPAD-502 readings as LCC = 1.26 x SPAD + 6.63. **(C)** Reflectance and **(D)** transmittance in the photosynthetically active radiation band (PAR, 400-700 nm), and **(E)** reflectance and **(F)** transmittance in the near-infrared band (NIR, 700-2500 nm), each averaged over the band from per-plot PROSPECT-D runs (Methods S1.3). B shows plots with a SPAD reading, and C-F show plots with a measured SPAD or leaf mass per area. Boxes show medians and interquartile ranges, and n is given above each box.

**Figure S4.**
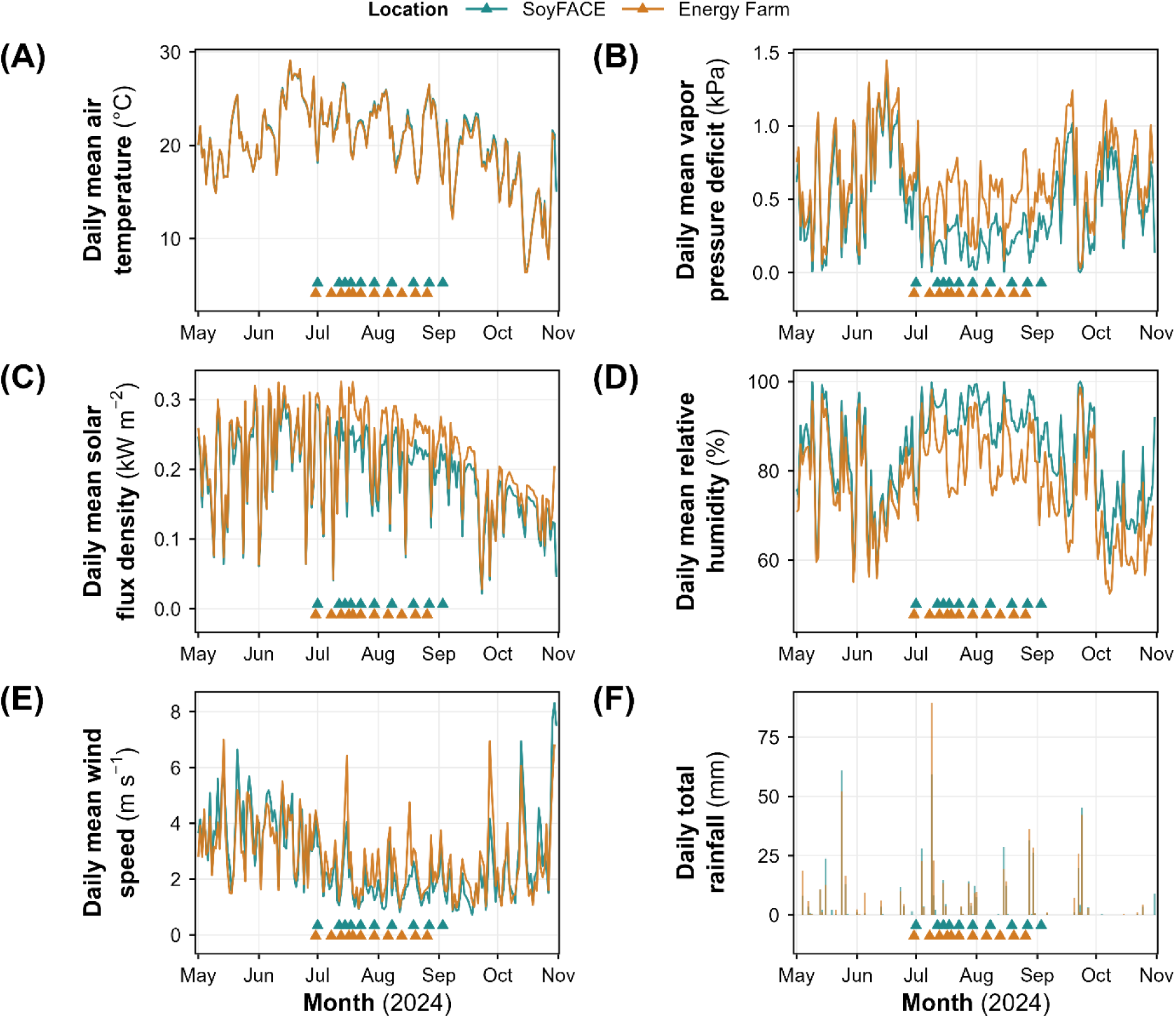
Daily weather at SoyFACE and the Energy Farm, Illinois, from 1 May to 31 October 2024. **(A)** Mean air temperature. **(B)** Mean vapor pressure deficit, computed from station vapor pressure and air temperature (Tetens equation) at the recording interval and then averaged by day. **(C)** Mean incoming solar flux density. **(D)** Mean relative humidity. **(E)** Mean wind speed. **(F)** Total rainfall. Data are hourly records from the SoyFACE station and 10**-**min records from the Energy Farm station. Triangles mark the dates on which leaf area index was measured (SoyFACE, upper row, 10 dates; Energy Farm, lower row, 11 dates).

**Figure S5.**
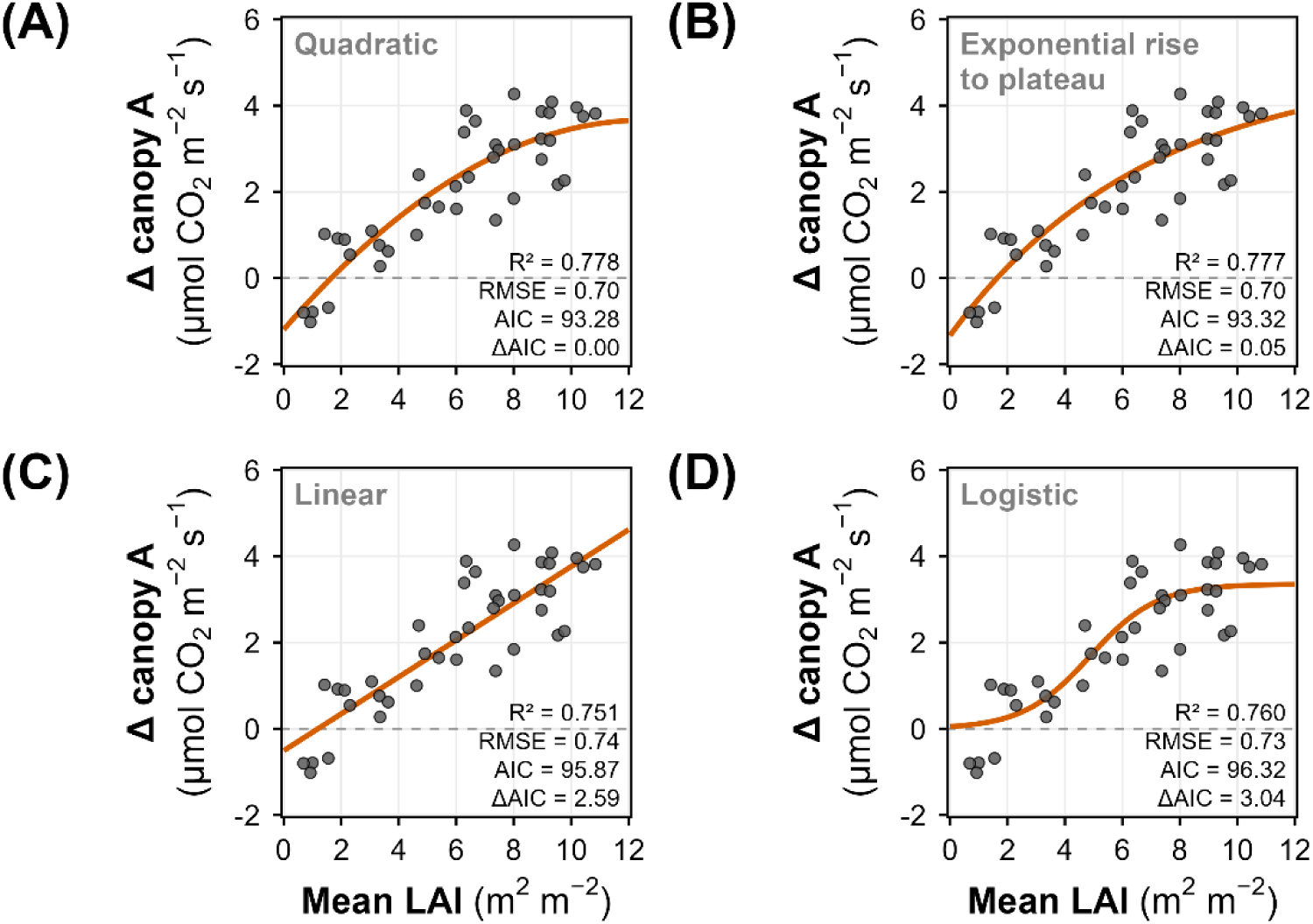
Alternative fits of the narrow-minus-broad difference in Helios canopy net photosynthesis (ΔA) against mean leaf area index (LAI) in broad- and narrow-leaflet soybean near-isogenic lines. Each panel shows the same 40 site x row spacing x date means with one fitted form. **(A)** Quadratic, ΔA = b₀ + b₁·LAI + b₂·LAI². **(B)** Exponential rise to a plateau, ΔA = c + a·(1 - exp(-k·LAI)). **(C)** Linear, ΔA = a + b·LAI. **(D)** Logistic, ΔA = a/(1 + exp(-k(LAI - L_half))). A and C were fitted by ordinary least squares and B and D by nonlinear least squares. Panels are ordered by Akaike information criterion (AIC), lowest first, and each gives R², the root-mean-square error (RMSE), AIC and the difference from the lowest AIC (ΔAIC). The dashed line marks ΔA = 0.

**Figure S6.**
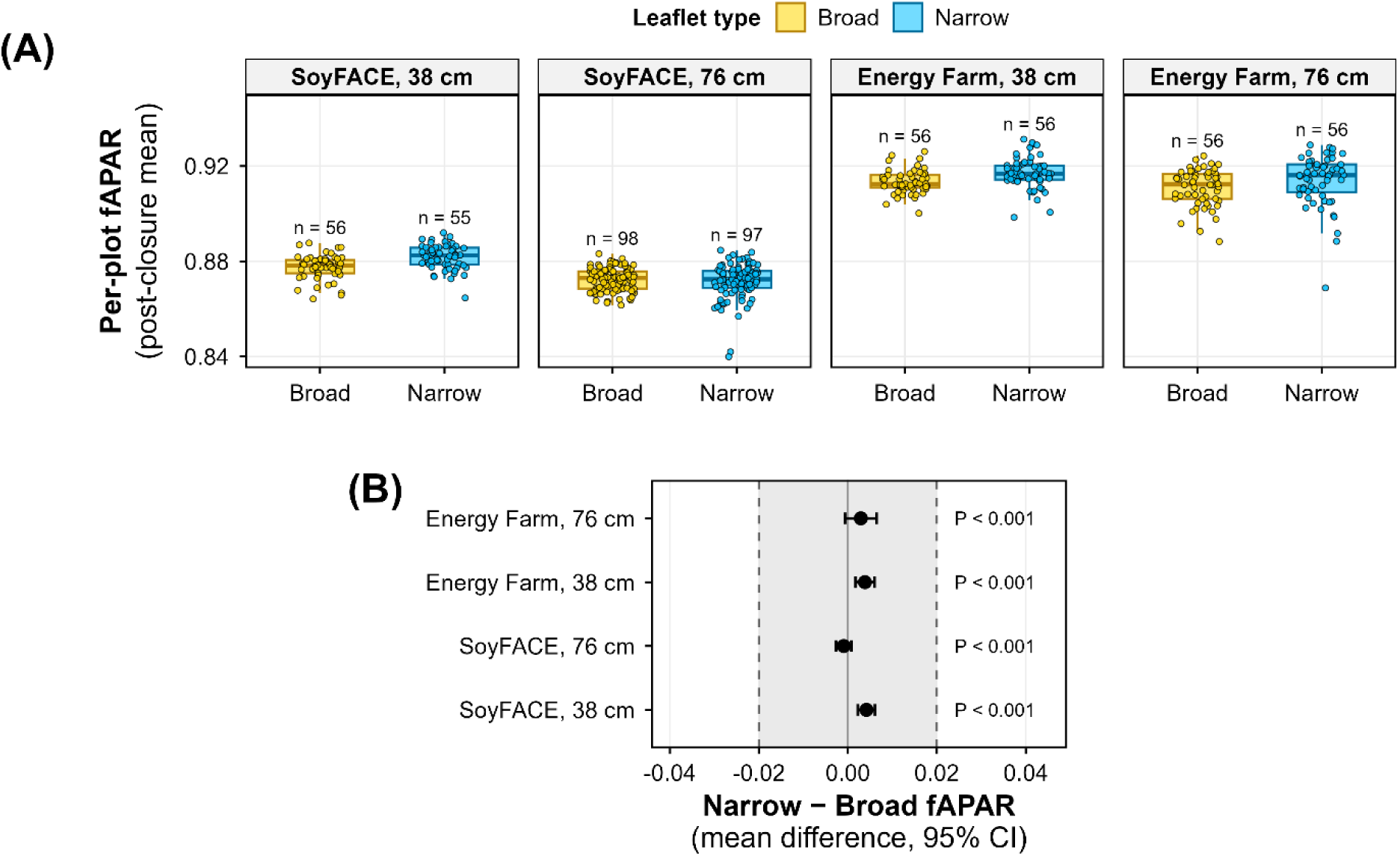
Fraction of incident photosynthetically active radiation absorbed (fAPAR) after canopy closure in broad- and narrow-leaflet soybean near-isogenic lines. **(A)** Per-plot fAPAR averaged over the period after canopy closure (≥ 70 days after planting) in each site x row spacing combination. Boxes show medians and interquartile ranges, and n is given above each box. **(B)** Difference between the narrow and broad leaflet-type means of fAPAR in each combination, with its 95% Welch confidence interval. The shaded band marks the ±0.02 equivalence margin. The P value for each row is from two one-sided tests (TOST), the larger of the two one-sided Welch t-test P values against the −0.02 and +0.02 margins.

**Figure S7.**
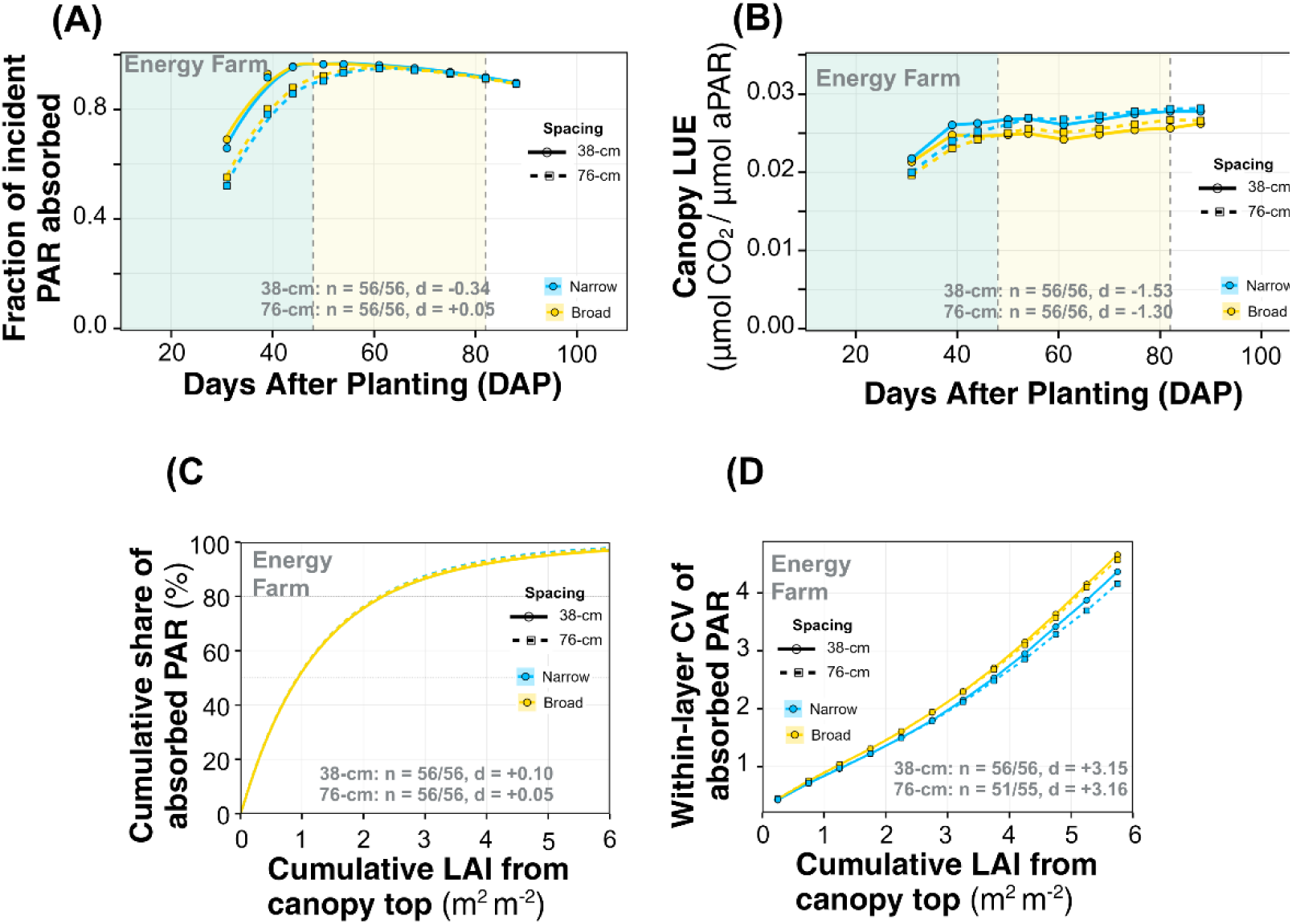
Energy Farm counterparts of Fig 3A-D for broad- and narrow-leaflet soybean near-isogenic lines. **(A)** Fraction of incident PAR absorbed (fAPAR) and **(B)** canopy light-use efficiency (LUE) against days after planting (DAP), as in Fig 3A and B. **(C)** Cumulative share of each plot’s absorbed PAR against cumulative leaf area index (LAI) from the canopy top, as in Fig 3C. **(D)** Leaf-area-weighted coefficient of variation (CV) of absorbed PAR among sub-leaf elements in successive 0.5-LAI layers, as in Fig 3D. C and D use each class’s peak-LAI date, which was DAP 61 (30 July 2024) for both leaflet types in 38-cm rows and DAP 68 (6 August, broad) and DAP 75 (13 August, narrow) in 76-cm rows. Panels give n (narrow/broad plots) and Cohen’s *d* as in Fig 3A-D.

**Figure S8.**
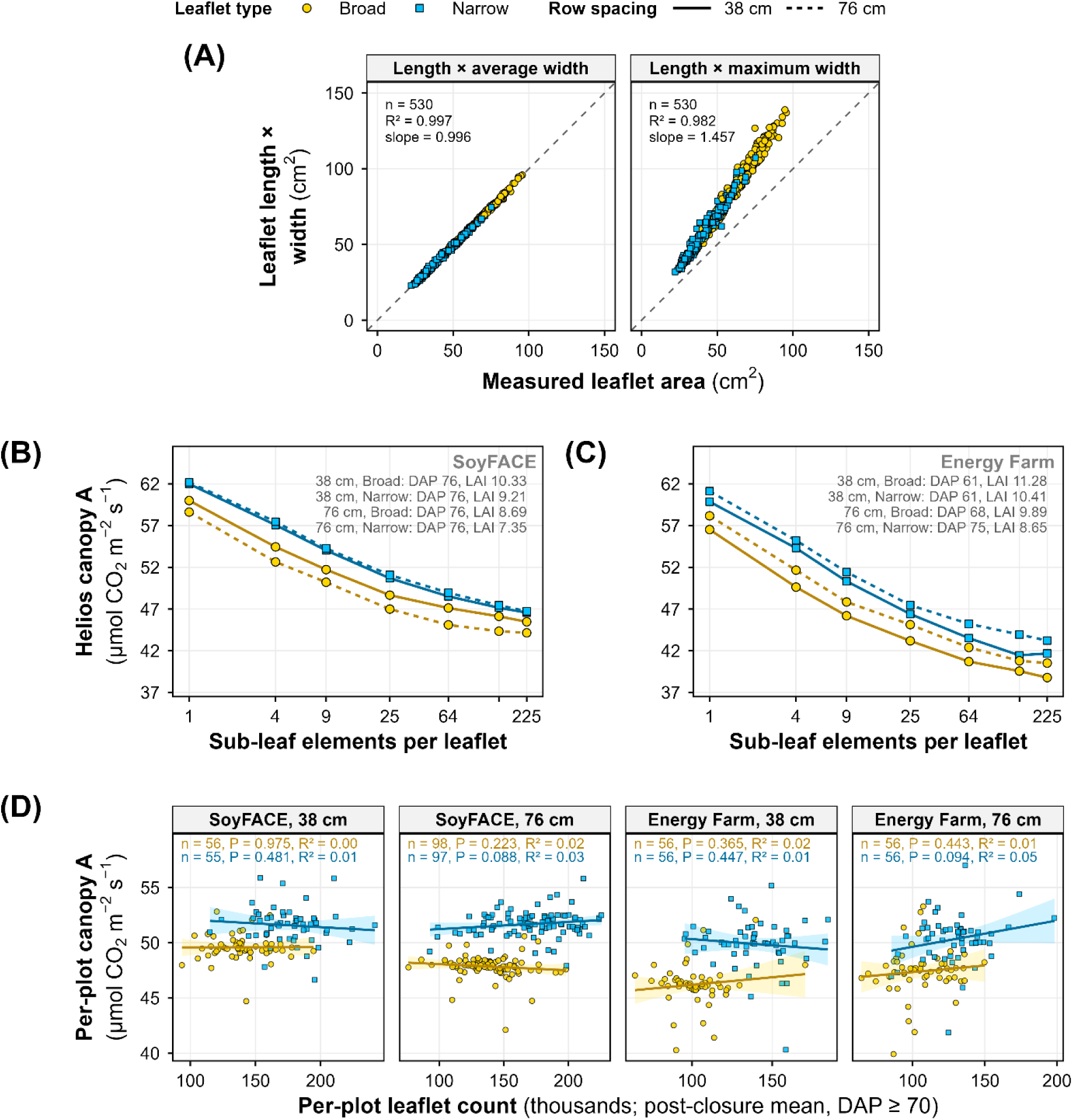
Checks on the Helios canopy simulations for broad- and narrow-leaflet soybean near-isogenic lines. **(A)** Leaflet length times average width (left) and times maximum width (right) against measured leaflet area for 530 plots, with the 1:1 line dashed and n, R² and slope in each panel. **(B-C)** Helios canopy net photosynthesis (canopy A) against the number of sub-leaf elements per leaflet (log scale) at SoyFACE (B) and the Energy Farm (C), for each leaflet type and row spacing on its peak leaf area index (LAI) date (dates and LAI in the panels). **(D)** Per-plot canopy A against the number of leaflets in the simulated canopy, both averaged after canopy closure (≥ 70 days after planting), in each site x row spacing combination. Lines and ribbons are ordinary least-squares fits with 95% confidence intervals, and each panel gives n, the slope P and R² for each leaflet type.

## Notes

https://doi.org/10.13012/B2IDB-9478251_V1)

### Competing Interest Statement

The authors have declared no competing interest.

